# H3K36 Methylation - a Guardian of Epigenome Integrity

**DOI:** 10.1101/2024.08.10.607446

**Authors:** Reinnier Padilla, Gerry A. Shipman, Cynthia Horth, Eric Bareke, Jacek Majewski

## Abstract

H3K36 methylation is emerging as a key epigenetic modification with strong implications in genetic disease and cancer. However, the mechanisms through which H3K36me impacts the epigenome and asserts its functional consequences are far from understood. Here, we use mouse mesenchymal stem cell lines with successive knockouts of the H3K36 methyltransferases: NSD1, NSD2, SETD2, NSD3, and ASH1L, which result in progressive depletion of H3K36me and its complete absence in quintuple knockout cells, to finely dissect the role of H3K36me2 in shaping the epigenome and transcriptome. We show that H3K36me2, which targets active enhancers, is important for maintaining enhancer activity, and its depletion results in downregulation of enhancer-dependent genes. We demonstrate the roles of H3K36me2/3 in preventing the invasion of gene bodies by the repressive H3K27me modifications. Finally, we observe a previously undescribed relationship between H3K36me and H3K9me3: Following the depletion of H3K36me2, H3K9me3 is redistributed away from large heterochromatic domains and towards euchromatin. This results in a drastic decompartmentalization of the genome, weakening the boundaries between active and inactive compartments, and a catastrophic loss of long-range inter-compartment interactions. By studying cells totally devoid of H3K36 methyltransferase activity, we uncover a broad range of crucial functions of H3K36me in maintaining epigenome integrity.

## Introduction

DNA is wrapped around histones to form nucleosomes, the structural and functional unit of chromatin. Chromatin can be densely packed as heterochromatin, or loosely packed as euchromatin. The extent of compaction, and thereby access to DNA, is in part regulated by chemical modifications to the histone tails, known as epigenetic or histone post-translational modifications (PTMs). These histone PTMs form a highly interdependent network, where euchromatic modifications facilitate access to DNA and heterochromatic modifications restrict access to DNA^1–3^. The different types of modifications at histone residues induce steric changes that alter nearby or interacting residues.

Methylation of histones plays pivotal roles in modulating transcriptional processes, primarily by influencing the accessibility of chromatin and serving as a substrate for a multitude of effector proteins to exert their functions. The lysine residues on the N-terminal tails of histones can receive up to three methylation marks, and the degree of methylation at specific residues is often associated with distinct transcriptional states and genomic features^4,5^. While the methylation states of certain residues coincide on the same histone tail and support similar processes, the methylation states of other residues are antagonistic, often due to steric bulk, and support opposing processes. In addition, the collective effects of histone methylation significantly influence the organization and 3D architecture of interphase chromatin^6–11^. Over the past decade, methylation of histone 3 lysine 36 (H3K36me) has increasingly emerged as a critical component of the epigenome. When aberrantly regulated, either through mutations affecting the genes of its writers or of histone 3 (H3) genes, developmental disorders or cancer ensues, underscoring its functional significance^12–18^. H3K36me exists in mono- (H3K36me1), di- (H3K36me2), or tri-methylated (H3K36me3) states, and there are five well-established enzymes capable of depositing H3K36me: SETD2, NSD1, NSD2, NSD3, and ASH1L. Briefly, SETD2 tethers to the elongating RNAPII complex within actively transcribed genes to deposit nearly all H3K36me3^19,20^. In contrast, the NSD family of lysine 36 methyltransferases (K36MTs) have established roles in the deposition of the lower methylation states, H3K36me1/2. NSD1 and NSD2 have been found to be responsible for the majority of global H3K36me2, in both IGRs and genes, and their individual contributions appear to vary by cell type and developmental stage^2,14,18,21–23^. In comparison, we have previously found that NSD3 has a more distinct role in the deposition of H3K36me2, primarily exerting its catalytic activity at active cis-regulatory elements (CREs) - most notably at enhancers and promoter flanking regions^22^. Finally, ASH1L is the most specific and least prolific of the K36MTs, primarily depositing H3K36me2 at specific developmentally important genes^22^.

Since H3K36me is generally associated with transcriptionally active, euchromatic regions of the genome, it is presumed to play a role in maintaining and fine-tuning transcription. It is one of the modifications broadly flanking CREs, whose regulatory states are frequently inferred by the enrichment profiles and co-localization of several histone modifications^24^. Notably, H3K4 methylation (H3K4me) and H3K27 acetylation (H3K27ac) are amongst the most well characterized modifications that distinguish active CREs. Functionally, H3K4me1 serves to fine tune enhancer activity by acting as a substrate for the binding of chromatin remodelers and other protein effectors^25^, while H3K27ac has a more direct effect on chromatin by neutralizing the positively charged histone tails, thereby weakening their interaction with DNA and establishing an environment permissive for transcription^26^. The co-presence of H3K4me1 and H3K27ac is frequently used to identify active enhancers^24,27^. In contrast, focal peaks of H3K4me3 and H3K27ac are hallmark chromatin signatures of active promoters^24,28^. H3K36me2 has been found to exist in broad domains at both active enhancers and promoters in most assayed cell types^2,18,21–23^. Loss of broad H3K36me2 domains, resulting from either pathological mutations or genetic ablation of its writer enzymes, has been shown to impact the distribution of H3K4me1/3 and H3K27ac, and is associated with altered gene expression^18,23,29^. While the general relationship between H3K36me2, CREs and gene expression has been documented, it has always been studied in the context of very broad, megabase scale domains, and the precise role of H3K36me2 in maintaining the activity of CREs and its influence on the deposition of other active histone marks in a localized setting has yet to be carefully examined.

Our understanding of the relationship between H3K36me and heterochromatin is also far from complete. On one hand, there exists a well documented antagonism between H3K36me2/3 and the silencing marks H3K27me2/3. The PRC2 complex, which deposits all H3K27me, is allosterically inhibited from nucleosomes marked with H3K36me2/3^30,31^. As a result, particularly H3K27me3 is excluded from H3K36me2/3 domains. While H3K27me2/3 are considered to be repressive marks, they are generally found in gene-rich regions and their presence is associated with facultative heterochromatin^32^. In comparison, H3K9me3 is distinctive of constitutive heterochromatin, and broad H3K9me3 domains generally blanket gene-poor regions containing tandem repeat elements, such as found at telomeres and pericentromeric regions^6,33,34^. While broad genomic domains of H3K9me3 and H3K36me2 tend to be mutually exclusive, the relationship between H3K36me and H3K9me is currently unclear, largely because evidence for mechanistic links between the two marks is lacking.

One of the challenges associated with investigating the functional role of H3K36me is the broad nature of its distribution, which is exacerbated by the number of enzymes capable of depositing these modifications. We have previously used CRISPR-Cas9 gene editing to establish clonal cell lines harboring individual or multiple knockouts (KOs) of the K36MTs, for the purpose of deconstructing their individual and combined contributions to the H3K36me landscape in C3H10T1/2 mouse mesenchymal stem cells (mMSCs)^22^. Here, we employ these cell lines to investigate the effects of H3K36me in more specific contexts: we examine focal peaks of NSD3- and ASH1L-mediated H3K36me2 present at active CREs, and their subsequent erasure, to assess their functional influence on the local epigenome and transcriptome. We provide further mechanistic insights into the ability of H3K36me2/3 to exclude H3K27me2/3 from actively transcribed genes, and the downstream impact on other relevant epigenetic modifications in these regions. Unexpectedly, we uncover a relationship between H3K36me, H3K9me3 and the 3D architecture of the genome that appears to be conserved in both mouse and human cells. Overall, the progressive loss of H3K36me resulting from the sequential deletion of the K36MTs provides a unique opportunity to assess the role of H3K36me in maintaining the active state of CREs, transcription and genomic compartmentalization.

## Results

### NSD3-mediated H3K36me2 maintains the active state of enhancers and their target genes

We have previously found that knocking out NSD1/2 (DKO) in mMSCs results in a near total depletion of broad intergenic H3K36me2^35^ (Fig. 1a). Further KO of SETD2 in the triple KO cells (TKO) results in the loss of nearly all genic H3K36me2 and H3K36me3^22^ (Fig. 1a). The remaining focal H3K36me2 regions, deposited by NSD3, are primarily targeted to active CREs^22^. Upon subsequent KO of NSD3 in the NSD1/2/3-SETD2 quadruple KO cells (QKO), these broad peaks of H3K36me2 disappear (Fig. 1a). To further explore the role of H3K36me2 in these confined regions, we examined promoters and enhancers that remain accessible, as defined by the presence of an ATAC-seq peak in both the TKO and QKO conditions. Having previously established biological replicates for each cell line, here we used ChIP-seq (H3K36me2, H3K27ac, H3K4me3) and CUT&RUN (H3K4me1, H3K27me3) to profile the relevant repressive and active histone modifications (Supplementary Fig. 1a), ATAC-seq for chromatin accessibility, whole genome bisulfite sequencing (WGBS) for DNA methylation (DNAme), and RNA-seq for transcriptional state to document the downstream changes following the depletion of H3K36me2 at CREs and their associated genes.

**Fig. 1.**
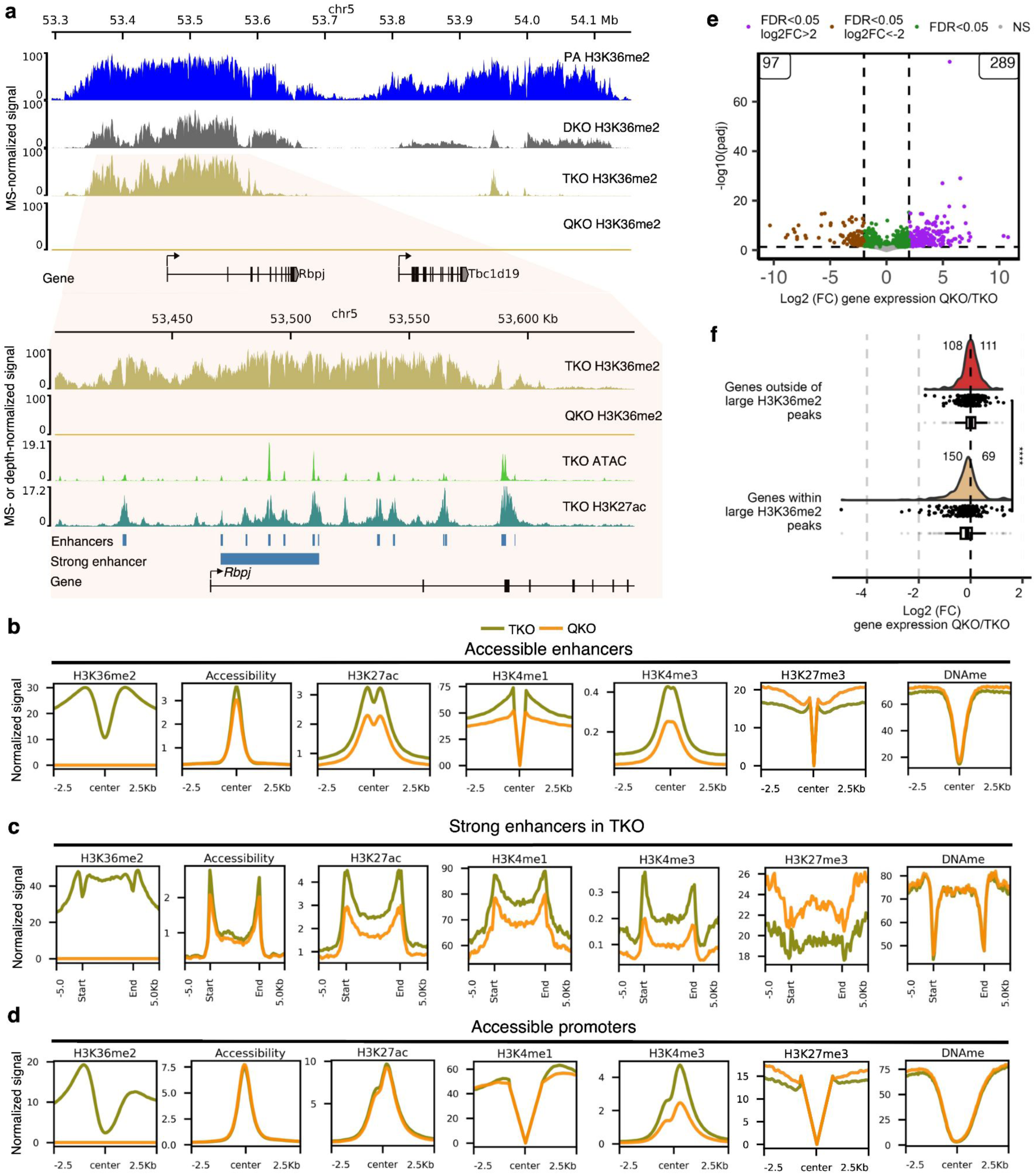
Loss of NSD3-dependent H3K36me2 leads to reduced enhancer activity and downregulation of targeted genes. **a** Representative genome browser tracks showing progressive loss of H3K36me2 in multiple-KO conditions: NSD1/2 double-KO (DKO); NSD1/2-SETD2 triple-KO (TKO); NSD1/2/3-SETD2 quadruple-KO (QKO). The zoomed-in tracks below illustrate that in TKO, the remaining NSD3-mediated H3K36me2 peaks mark several active and strong enhancers (dense clusters of nearby enhancers) that are lost in QKO. **b** Aggregate plots comparing TKO to QKO, indicating reduced chromatin accessibility, decreased signal of active marks (H3K27ac, H3K4me1, H3K4me3) and invasion of H3K27me3 at accessible enhancers (n=22706). **c** Aggregate plots centered on strong enhancers - defined as clusters of enhancers within 12.5 kb of each other (n=697) - showing further reduced activity for active marks (H3K27ac, H3K4me1 and H3K4me3) and more pronounced invasion of H3K27me3 following loss of H3K36me2. **d** Aggregate plots centered on accessible promoters showing no changes in accessibility, DNAme or H3K27ac, whereas H3K4me1/3 decrease and H3K27me3 increases comparing TKO to QKO (n=9599). **e** Volcano plot of differentially expressed genes, showing more genes become upregulated (289) than downregulated (97) comparing QKO to TKO. **f** Log2 fold-change plots of expression-matched genes within and outside of large H3K36me2 peaks (see **a** for example peak), demonstrating more genes becoming downregulated (150) than upregulated (69). An equal number of genes outside of these large H3K36me2 peaks were randomized and selected as control. Statistical significance was tested using the Wilcoxson signed rank test. **** represents *p*-value=5.8e-07. For **a, b, c,** and **d**, biological replicates (n=3) were merged. ATAC-seq tracks were depth-normalized (in counts per million (CPM)). For all other tracks, except for DNAme (which indicates the percent methylation at CpG sites), normalization factors were computed by multiplying genome-wide modification percentage values (averaged per condition) from mass-spectrometry (MS) by the total number of bins and dividing by the total signal for a given coverage track. This normalization factor was multiplied to the depth-normalized signal (CPM) for each track to generate MS-normalized signals. MS-normalized signals represent the mean local frequency of the relevant modification.

**Supplementary Figure 1.**
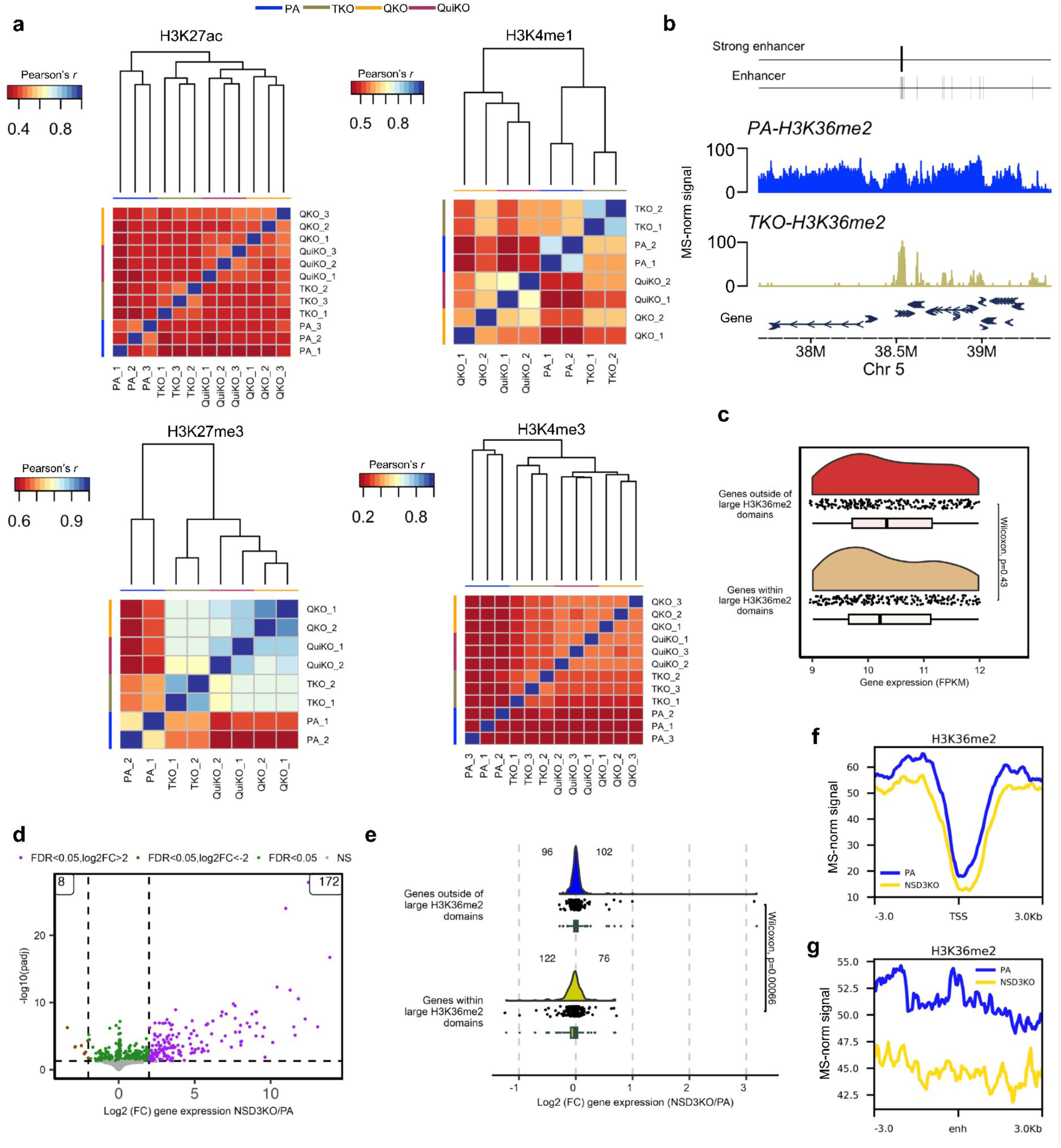
Analysis of the downstream consequences of focal loss of H3K36me2. **a** Pearson correlation heatmaps for various histone marks demonstrating that replicates cluster together. **b.** Representative genome-browser tracks showing NSD1/2-SETD2-TKO leads to depletion of broad H3K36me2 domains, with the remaining, smaller H3K36me2 regions (i.e. peaks) found at clusters of enhancers. **c.** Gene expression plots showing that the two gene groups (genes outside of large H3K36me2 peaks and genes within large H3K36me2 peaks) have no significant difference in basal expression (FPKM) in the parental (wildtype) cell line. **d.** Volcano plot showing differential gene expression between NSD3KO and parental cell lines. **e.** Gene expression log2 fold changes of NSD3KO versus parental cells showing that in the same set of genes within the large H3K36me2 peaks from TKO, 122 genes become downregulated whereas 76 become upregulated. In a control set of genes, a similar number of genes become upregulated (96) and downregulated (102). **f.** Aggregate plot of H3K36me2 centered on the TSS of the set of genes within large H3K36me2 peaks from **e,** showing a decrease in NSD3KO compared to parental samples. **g**. Aggregate plot of H3K36me2 signal centered on annotated enhancers closest to the genes from **f**, indicating a stronger decrease at the enhancers of these target genes. For **b, f** and **g**, MS-normalized signals were computed as previously described in Fig. 1, and represent the mean local frequency of the relevant modification.

At active enhancers, the depletion of H3K36me2 is accompanied by slight decreases in chromatin accessibility (Fig. 1b). Concurrently, we also observe a decrease in the levels of other known active enhancer modifications, specifically H3K4me1/3 and H3K27ac (Fig. 1b). In contrast, the levels of H3K27me3 increase at enhancers that are depleted of H3K36me2 (Fig. 1b), which was expected given the known antagonistic relationship between these marks^30,31^. Previous studies have shown that H3K36me2/3 serve as templates for the deposition of de novo DNAme by DNMT3A/B, respectively^35,36^. Therefore, we expected that in the absence of H3K36me2/3, enhancers may experience substantial reductions in CpG methylation. Using WGBS, however, we find that DNAme is largely unchanged (Fig. 1b). We also interrogated strong enhancers, as defined by dense clusters of individual enhancers (Fig. 1a; Supplementary Fig. 1b), which are known to be essential for the maintenance and regulation of genes crucial to both normal development and cancer^37^. In TKO cells, most strong enhancers retain H3K36me2, which is subsequently depleted in the QKO condition, demonstrating that NSD3 is recruited to these regions (Fig. 1a, c). Here, we found similar yet more pronounced trends in the decrease of activating and increase of silencing modifications, further demonstrating that H3K36me2 is essential to the maintenance of enhancer activity (Fig. 1c).

Interestingly, the changes occurring at the promoters of NSD3 target genes appear to be considerably more attenuated, as compared to those at enhancers. The overall levels of NSD3-deposited H3K36me2 in TKO cells are lower at accessible promoters than at accessible enhancers (Fig. 1b, c, d). Thus, while the loss of H3K36me2 is nearly complete at all CREs, the relative decrease is smaller at promoters. Consequently, at promoters we observe no change to chromatin accessibility and H3K27ac, while the decrease of H3K4me1 and increase of H3K27me3 is lower than at enhancers (Fig. 1d). We do, however, observe a comparable 2-fold decrease in both promoter- and enhancer-marked H3K4me3 (Fig. 1d). Overall, these results provide further evidence that the presence of NSD3-associated H3K36me2 is particularly influential at enhancers.

We next examined the effects on gene expression following the depletion of CRE-associated H3K36me2. Genome-wide, we found no general trend towards downregulation, with more than twice as many genes going up in expression (289) than down (97) (Fig. 1e). However, many of these changes may be secondary, downstream effects from global H3K36me2 depletion and/or ablation of NSD3. In view of the more pronounced epigenetic changes occurring at enhancers, we hypothesized that the most direct effect on gene expression should be observed at genes whose expression is dependent on H3K36me2-marked enhancers. We identified a set of predicted H3K36me2-dependent genes based on their overlap with large H3K36me2 peaks that contained at least one annotated enhancer (from Ensembl and FANTOM databases). We also created a complimentary, randomized control set of H3K36me2-independent genes. The two sets were further matched for baseline gene expression levels by selecting only transcripts within the 9-12 FPKM range (Supplementary Fig. 1c). We found that following the loss of NSD3-dependent H3K36me2, genes associated with large H3K36me2 peaks become significantly more downregulated (150 down versus 69 up) compared to the control set of genes (108 down versus 111 up) (Fig. 1f). Our findings indicate that the depletion of NSD3-dependent H3K36me2 impacts gene expression primarily by reducing enhancer activation of target genes rather than directly influencing promoter activity.

We have previously examined the effects of the individual KO of NSD3 on the global H3K36me2 landscape, and we did not observe a significant depletion on the global abundance of H3K36me2, possibly because of compensation by the other K36MTs^22^. To investigate whether the effects on CREs are a result of the H3K36me2 catalytic product or driven by NSD3 itself, we analyzed the same set of enhancer-dependent NSD3 target genes identified in the TKO cells in the context of comparing parental to NSD3-KO cells. Although globally, we find that many more genes significantly increase in expression (172) than decrease (8) (Supplementary Fig. 1d), within the set of predicted H3K36me2-dependent genes derived from the TKO cells, we found a tendency towards downregulation (122) versus upregulation (76) (Supplementary Fig. 1e). This suggests that the compensation of H3K36me2 levels by the other K36MTs is not complete. Upon closer inspection, we observe a decrease of H3K36me2 at the promoters of this set of genes in NSD3-KO cells (Supplementary Fig. 1f) and an even greater decrease at the nearest enhancers of these promoters (Supplementary Fig. 1g). Hence, even in parental cells, where NSD3 is not the dominant K36MT, its absence is reflected in a slight, localized reduction of H3K36me2 and downregulation of enhancer-dependent genes.

Overall, our results support that while NSD3-mediated H3K36me2 is targeted to both enhancers and promoters, it is most influential at enhancers with regards to maintaining their active chromatin states, restricting the invasion of H3K27me3, and has discernible effects on maintaining the expression of their target genes.

### Functional characterization of focal H3K36me2 at ASH1L target genes

In the QKO cells, we had previously identified 119 remaining H3K36me2 peaks, with the majority of these peaks straddling the transcription start sites (TSS) (n=60) or located within the gene bodies (n=24) of a subset of developmentally important genes^22^. Those peaks disappeared in the NSD1/2/3-SETD2-ASH1L-QuiKO (QuiKO) condition, following the ablation of ASH1L (Fig. 2a). Similar to the comparison between TKO and QKO cells, this provides a unique opportunity to study the functional consequences of H3K36me2 depletion in a very specific as opposed to a global setting. We find that the majority of ASH1L target genes (62/84, *p* = 0.0015, Wilcoxon test) decrease in expression following the loss of ASH1L (Fig. 2b, c). Furthermore, the change in expression is proportional to the size of the original H3K36me2 peaks: genes with the largest H3K36me2 peaks in QKO cells have the largest fold changes in gene expression (Fig. 2d). In contrast, the same set of genes did not decrease in expression when comparing TKO to QKO cells, where H3K36me2 is unaffected at these promoters (Fig. 2b, c). Finally, it is possible that ASH1L itself may act as a transcriptional modulator, and may have an effect independent of its catalytic activity - as has been recently suggested in the case of NSD1^23^. To distinguish whether the reduced expression of the 84 ASH1L-target genes may be due to the loss of ASH1L itself, or its catalytic product, H3K36me2, we looked at differences in expression between parental mMSCs and the individual ASH1L-KO cells. In the ASH1L-KO condition, the levels of H3K36me2 remain largely unchanged at the promoters of these genes, most likely as a result of compensation by the other K36MTs, which exhibit less specificity in their deposition of H3K36me2 (Supplementary Fig. 2a). Interestingly, we found no significant changes to the expression of ASH1L target genes, suggesting that the presence of H3K36me2, rather than the presence of ASH1L has a role in maintaining their expression (Fig. 2c).

**Fig. 2.**
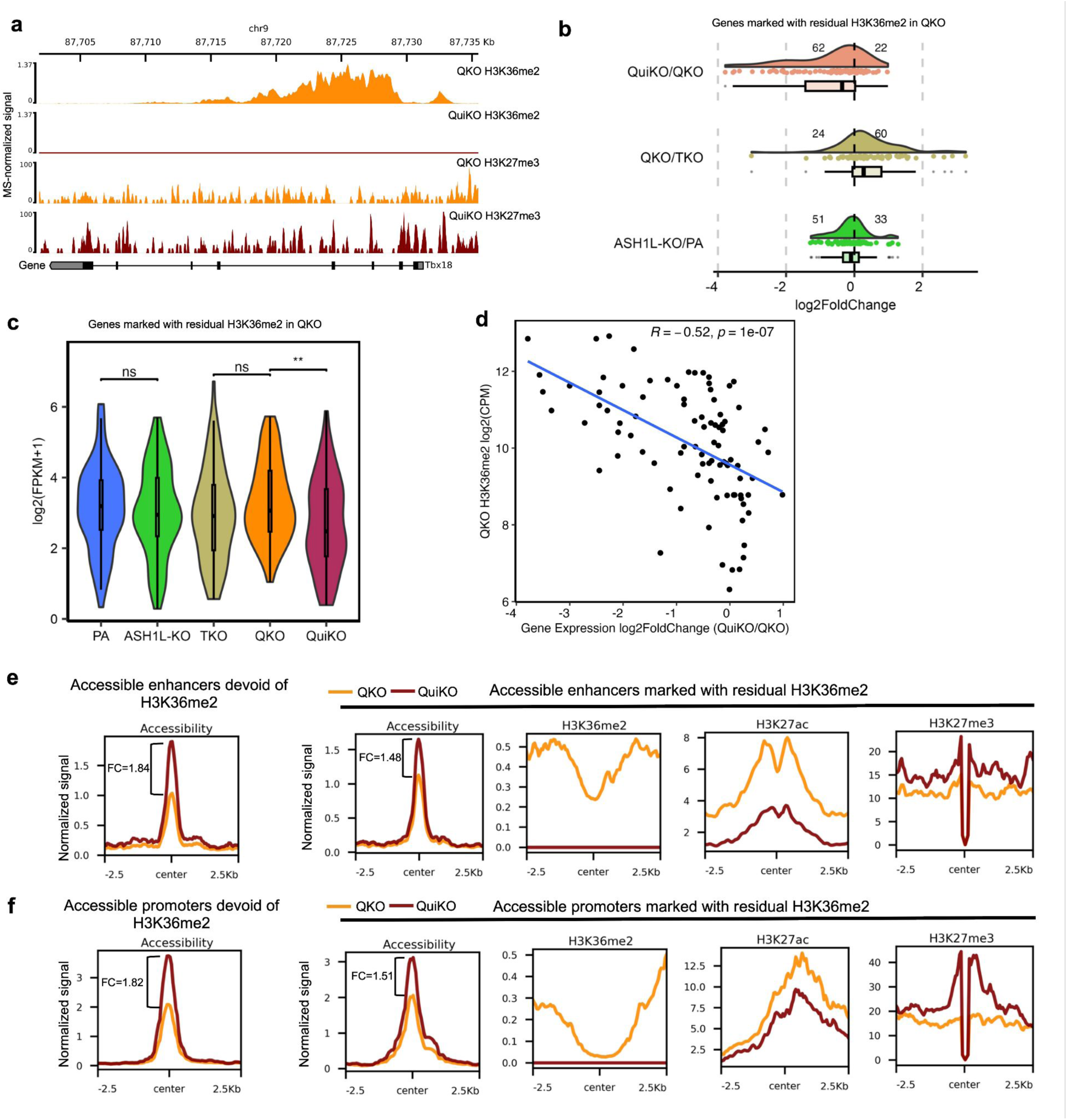
Loss of ASH1L-mediated H3K36me2 results in the downregulation of genes marked by H3K36me2, decreased H3K27ac and increased H3K27me3 at cis-regulatory elements. **a** Genome-browser tracks showing one of 84 genes that is still marked by residual H3K36me2 in QKO cells, all of which is lost following ASH1L-KO in QuiKO cells. **b** Log2 fold changes of gene expression for the 84 genes marked by residual H3K36me2 in QKO, showing that more genes become downregulated (62) than upregulated (22) following loss of ASH1L-mediated H3K36me2. Within this gene set, 51 genes are downregulated while 33 are upregulated when comparing ASH1L-KO to the parental cells. This trend of downregulation is not observed when comparing QKO to TKO for this gene set, where 60 genes are upregulated and only 24 are downregulated. **c** Violin plots showing that the 84 genes in **b** only significantly decrease in expression when comparing QKO to QuiKO. ** represents *p*-value <= 0.01 from Wilcoxson signed rank test. **d** Scatter plot illustrating a significant negative correlation between the size of the H3K36me2 peak and gene expression log2 fold changes following ASH1L-dependent H3K36me2 loss in the QKO cell line. Reported values are Pearson’s correlation coefficient (*R*) and the associated *p*-value. Each point represents a gene. **e-f** Aggregate plots showing H3K36me2 and H3K27ac decreases whereas H3K27me3 increases at accessible enhancers (n=72) and promoters (n=55) marked with residual H3K36me2. Genome-wide, chromatin accessibility increases in QuiKO cells compared to QKO, although the CREs marked with residual H3K36me2 gain considerably less. Greater decreases are found at enhancers than promoters marked with residual H3K36me2. For **f**, since an intersection of ATAC-seq peaks between QKO and QuiKO was used to identify accessible promoters, only 55 out of the 60 H3K36me2-flanked promoters had an ATAC-seq peak detected at their promoter. Normalized signals were either depth-normalized (ATAC-seq) or MS-normalized (all other tracks), as previously described. In the box plots for **b** and **c**, boxes span the lower (first quartile) and upper quartiles (third quartile), median is indicated with a center line and whiskers extend to a maximum of 1.5 times the interquartile range.

**Supplementary Figure 2.**
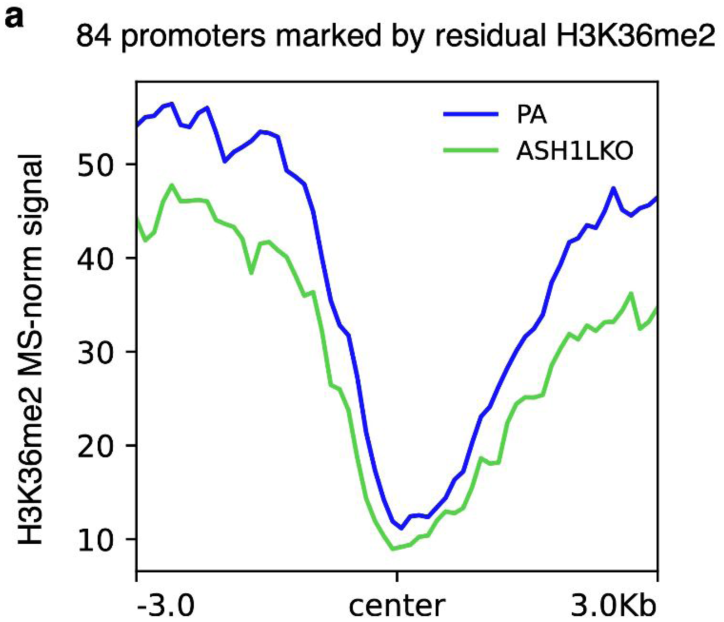
KO of ASH1L independently does not lead to significant depletion of H3K36me2. **a** Aggregate plot of MS-normalized H3K36me2 signal centered on the 84 promoters marked with residual H3K36me2 in the QKO cells, showing similar H3K36me2 levels between PA and ASH1L-KO at these promoters.

Concurrently, we investigated changes to other epigenetic modifications at the remaining CREs marked with residual H3K36me2. Genome-wide, we found that chromatin accessibility increases in QuiKO cells compared to QKO, however, the CREs marked with residual H3K36me2 gain considerably less (Fig. 2e, f). This likely reflects the net negative effect of losing H3K36me2 at these specific loci in comparison to other genomic regions. Nevertheless, the depletion of ASH1L-mediated H3K36me2 is accompanied by a further decrease of H3K27ac and a corresponding increase of H3K27me3 at these CREs (Fig. 2e, f). Similar to the loss of NSD3 in QKO cells, we observe that the loss of H3K27ac is greater at enhancers than promoters (Fig. 2e, f), further supporting that the presence of H3K36me2 at CREs may be more influential at enhancers than promoters. Altogether, these analyses demonstrate that H3K36me2 contributes to maintaining the balance between activating and silencing epigenetic modifications at CREs, and this has consequences for the expression of target genes.

### The effect of H3K36 methylation on antagonizing H3K27me within genes

One of the presumed functions of H3K36me3 that is deposited within actively transcribed genes is to prevent the deposition of the silencing modifications at H3K27^30,31^. This effect may further be strengthened by the presence of H3K36me1/2. It has been shown that methylation at H3K36 antagonizes PRC2 and hinders the deposition of methylation at H3K27^30,31,38^. This is particularly true for the highest methylation levels, resulting in mutual exclusivity of H3K36me3 and H3K27me3 on the same histone tail^31,38^. Accordingly, H3K27me3 is excluded from actively transcribed genes, H3K27me2 is generally depleted, and only H3K27me1 invades gene bodies, where its distribution is similar to that of H3K36me3 (Fig. 3a).

**Fig. 3.**
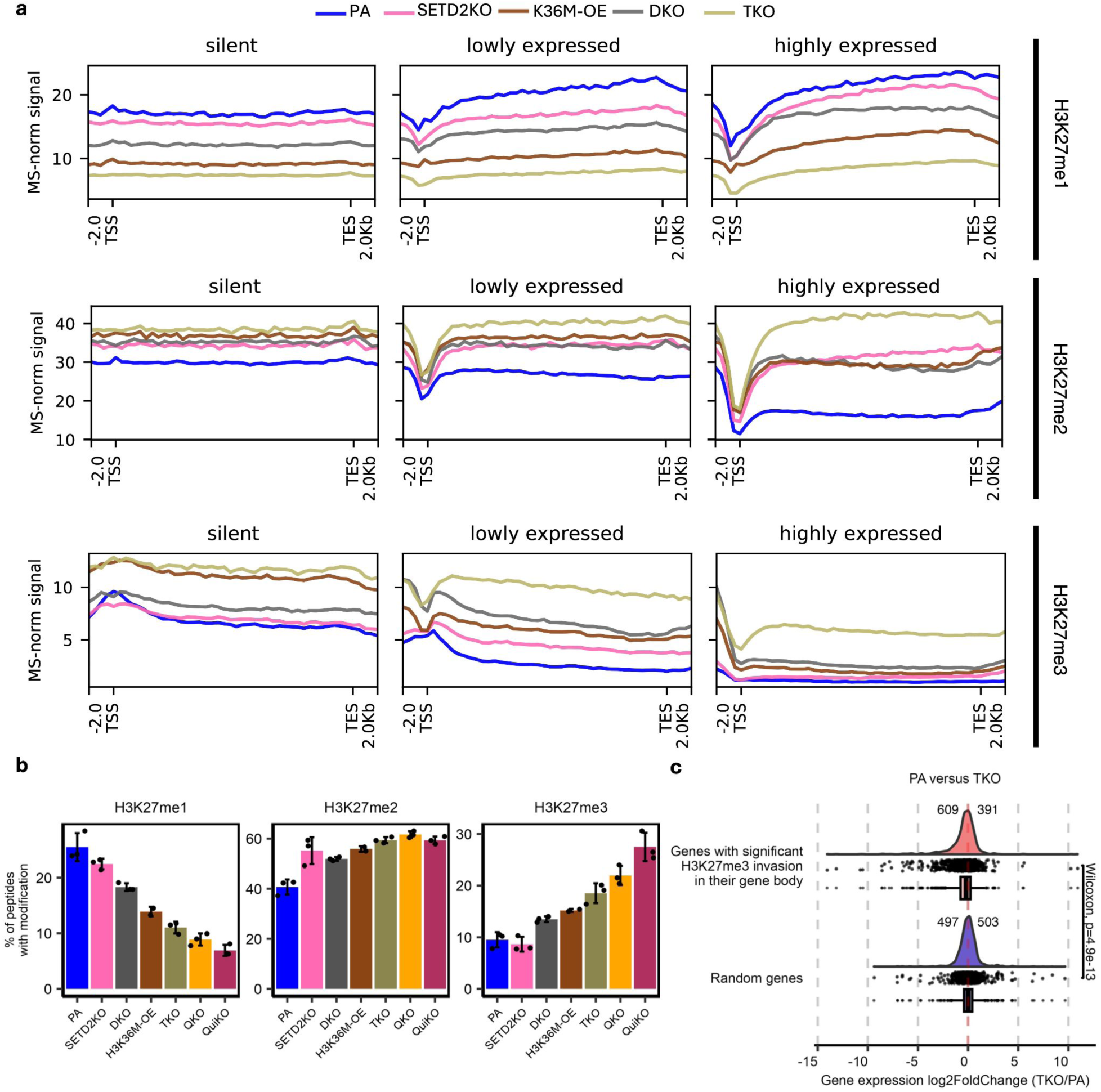
Invasion of H3K27me3 into gene bodies following depletion of H3K36me. **a** Metagene plots depicting the invasion of H3K27me within gene bodies. Protein-coding genes with gene lengths falling between the 25th and 75th percentiles and with FPKM values exceeding 0.1 were included. After filtering and stratifying into three quantiles, the bottom 2000 were classified as lowly expressed and the top 2000 as highly expressed genes. 2000 genes with zero expression were designated as silent genes. ChIP-seq H3K27me1/2/3 signals were MS-normalized as previously described. **b** Barplots of genome-wide prevalence of modifications based on mass spectrometry, showing progressive depletion of H3K27me1, and progressive increases in H3K27me2 and H3K27me3 levels in multi-KO conditions reaching approximately 1.5-fold and more than 2.5-fold respectively. Error bars display the standard deviation around the mean. (n=3 per condition). **c** Log2 fold changes of gene expression comparing PA to TKO, depicting 1.5-fold more genes being downregulated (609) than upregulated (391) following significant H3K27me3 invasion (adjusted *p*-value < 0.05) into their gene bodies compared to a randomized, control set of genes, which have similar numbers of genes up- (503) and down-regulated (497). Statistical significance was derived from Wilcoxson signed rank test. In the box plots, boxes span the lower (first quartile) and upper quartiles (third quartile), median is indicated with a center line and whiskers extend to a maximum of 1.5 times the interquartile range.

**Supplementary Figure 3.**
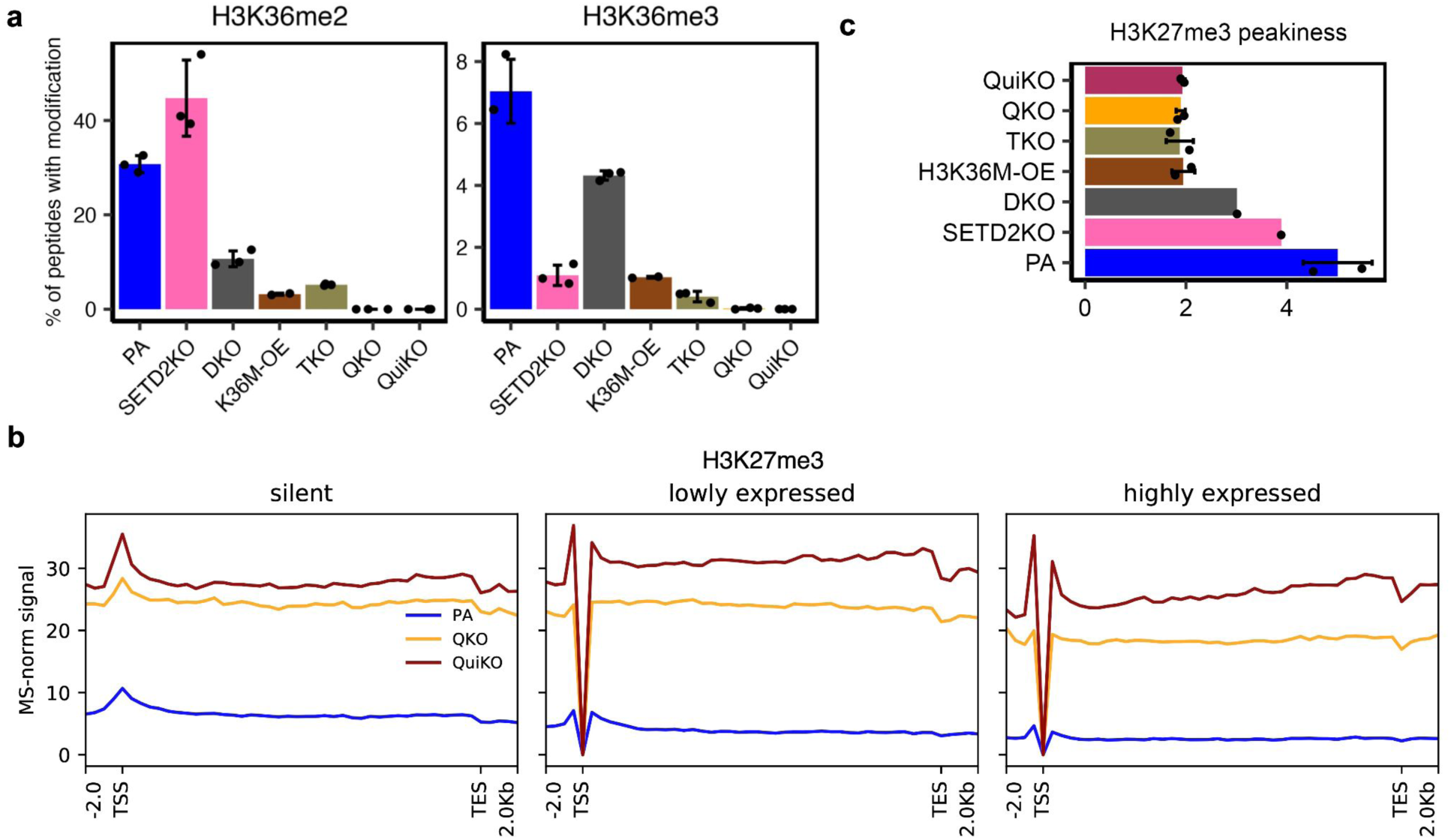
Analysis of H3K27me3 spreading following H3K36me depletion. **a** Barplots of genome-wide prevalence of modifications based on mass spectrometry showing global loss of H3K36me2/3 following knockouts of K36MTs. **b** Metagene plots of MS-normalized H3K27me3 CUT & RUN signal plotted within the same sets of genes from Fig. 3a, showing increasing invasion of H3K27me3 into gene bodies with further depletion of H3K36me2 in QKO and QuiKO. **c** H3K27me3 “peakiness” plot showing decreased “peakiness” scores, indicating spreading of H3K27me3 (broader distribution) following multiple knockouts of K36MTs. For **a** and **c**, error bars display the standard deviation around the mean. (n=3 per condition).

Using our panel of CRISPR engineered mMSCs, we investigated the effects of H3K36me depletion on the presence of H3K27me in gene bodies. We used quantitative mass spectrometry (MS) to quantify the changes to the absolute abundance of each of the respective methylation marks, and ChIP-seq/CUT&RUN to profile changes to the global distributions of H3K27me states.

Unexpectedly, in the SETD2 mutant which lacks nearly all H3K36me3 (Supplementary Fig. 3a), the increase of H3K27me3 in gene bodies is minimal, and primarily discernible within lowly transcribed genes (Fig. 3a; bottom row, middle box). However, the absence of H3K36me3 has a more pronounced effect on H3K27me2 and permits the deposition of this mark within genes with only a weak dependence on transcription and a corresponding decrease in H3K27me1 (Fig. 3a; middle row). Accordingly, these genic changes are reflected in the overall genome-wide abundance: H3K27me2 slightly increases and this is accompanied by a proportional decrease of H3K27me1, while changes to H3K27me3 are minimal (Fig. 3b). These observations may be explained by the fact that the sole loss of SETD2 in the presence of the other intact K36MTs primarily depletes H3K36me3, and results in an increase of genic H3K36me2 due to fewer residues being upgraded to the higher methylation state^22^ (Supplementary Fig. 3a). Since the presence of H3K36me2 also hinders the activity of PRC2 to deposit higher orders of H3K27me^30^, these results suggest that the presence of H3K36me3 inhibits the deposition of both H3K27me2/3, while the presence of H3K36me2 may primarily inhibit the deposition of H3K27me3.

Therefore, we next focused specifically on the ability of H3K36me2 to protect gene bodies from methylation at H3K27. In NSD1/2-DKO cells, the levels of H3K36me2 are significantly reduced, with a moderate reduction of H3K36me3 (Supplementary Fig. 3a). This is accompanied by a significant increase in the global levels of H3K27me2/3 (Fig. 3b). Interestingly, the partial reduction of H3K36me2 has a more pronounced effect on genic H3K27me2/3 than what is observed with the near total depletion of H3K36me3 (Fig. 3a; middle and bottom rows). Although in vitro studies indicate a stronger antagonistic effect from H3K36me3^30^, the greater observable effect of H3K36me2 depletion may reflect the generally higher absolute levels of H3K36me2 than H3K36me3 in gene bodies. Finally, we used NSD1/2-SETD2-TKO cells to study the consequences of depleting both H3K36me2/3. Expectedly, in these cells we observe the strongest invasion of H3K27me2/3 into gene bodies (Fig. 3a; middle and bottom rows). Both H3K27me2/3 now appear free to populate genes, with little dependence on expression levels (Fig. 3a). Further depletion of H3K36me2 in our QKO and QuiKO cell lines leads to even greater invasion of H3K27me3 into gene bodies (Supplementary Fig. 3b). Interestingly, the cells overexpressing the H3K36M oncohistone (H3K36M-OE), which reduces both H3K36me2/3, show invasion of H3K27me that is intermediate between the DKO and TKO conditions (Fig. 3a, b).

To better illustrate the global changes to the distribution of H3K27me across the various conditions, we calculated H3K27me3 ‘peakiness scores’, which represents the average ChIP-seq/CUT&RUN signal in the top 1% of 1-kb bins over the total signal, confirming the spreading (lower peakiness score) of H3K27me3 following sequential removal of H3K36me (Supplementary Fig. 3c). The lower peakiness scores in the multiple-KO conditions demonstrates that as H3K36me is sequentially removed, the distribution of H3K27me3 shifts from a focused distribution in parental cells, to a broader distribution in cells lacking H3K36me. Lastly, more than 1.5-fold of genes with significant H3K27me3 invasion in their gene body decrease in expression (609 down-compared to 391 upregulated), further supporting that hypothesis that the invasion of H3K27me3 into gene bodies contributes to suppressing gene expression (Fig. 3c).

Overall, while our results demonstrate that H3K36me plays a role in excluding H3K27me2/3 from gene bodies, additional factors are clearly responsible for preventing the full invasion of H3K27me. The two likely culprits are 1) nucleosome turnover, which results in replacement of modified with naïve nucleosomes during transcription^39^, and 2) DNAme, which may hinder the recognition of CpG rich sequences, such as those found in exons, by PRC2^40^.

### The effect of H3K36 methylation on other epigenetic modifications within gene bodies

As previously described, H3K36me2/3 have been shown to serve as substrates for the localization of DNMT3A/B to deposit de novo DNAme^35^. Therefore, we expected that in the absence of H3K36me2/3, gene bodies may experience substantial reductions in CpGme. However, we find that even in the QuiKO cells, which lack nearly all H3K36me, the most highly expressed genes retain DNAme levels comparable to parental cells (Fig. 4a; third box). Methylation is visibly reduced in the lowest expression quantile, as well as in unexpressed genes – where the latter behave similarly to inactive intergenic regions where DNA hypomethylation has previously been shown to be associated with the reduction of intergenic H3K36me2 (Fig. 4a; first and second box)^2,18,35^. It appears, however, that in highly expressed genes, H3K36me-mediated targeting of de novo DNMTs is unnecessary for the maintenance of high DNAme levels.

**Fig. 4.**
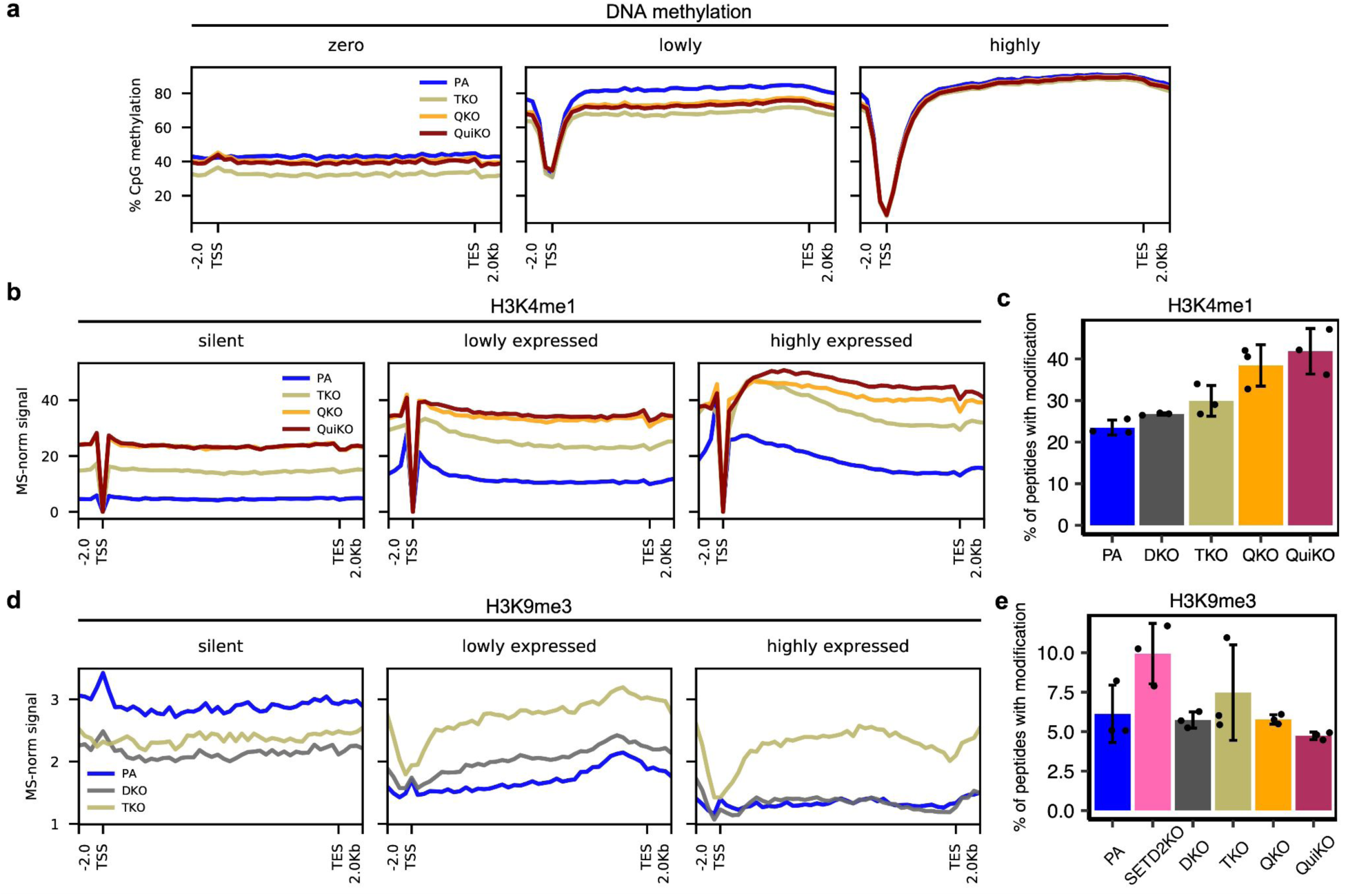
Depletion of H3K36me induces changes in DNA methylation, H3K4me1 and H3K9me3 within gene bodies. **a** Metagene plots of DNA methylation within gene bodies (same groups of genes used in Fig. 3), illustrating a decrease in DNA methylation in unexpressed and lowly expressed genes. **b** Metagene plots indicating increasing H3K4me1 within gene bodies following depletion of H3K36me. **c** Barplot illustrating the increasing global abundance of H3K4me1, as determined by mass spectrometry (MS), following successive deletions of K36MTs. **d** Metagene plots indicating invasion of H3K9me3 within gene bodies following depletion of H3K36me2 in DKO and TKO cell lines. **e** Barplot derived from MS, showing that genome-wide H3K9me3 abundance does not significantly change across KO cell lines. For **b** and **d**, CUT&RUN or ChIP-seq signals, respectively, were MS-normalized as previously described. For **c** and **e**, error bars display the standard deviation around the mean. (n=3 per condition).

**Supplementary Figure 4.**
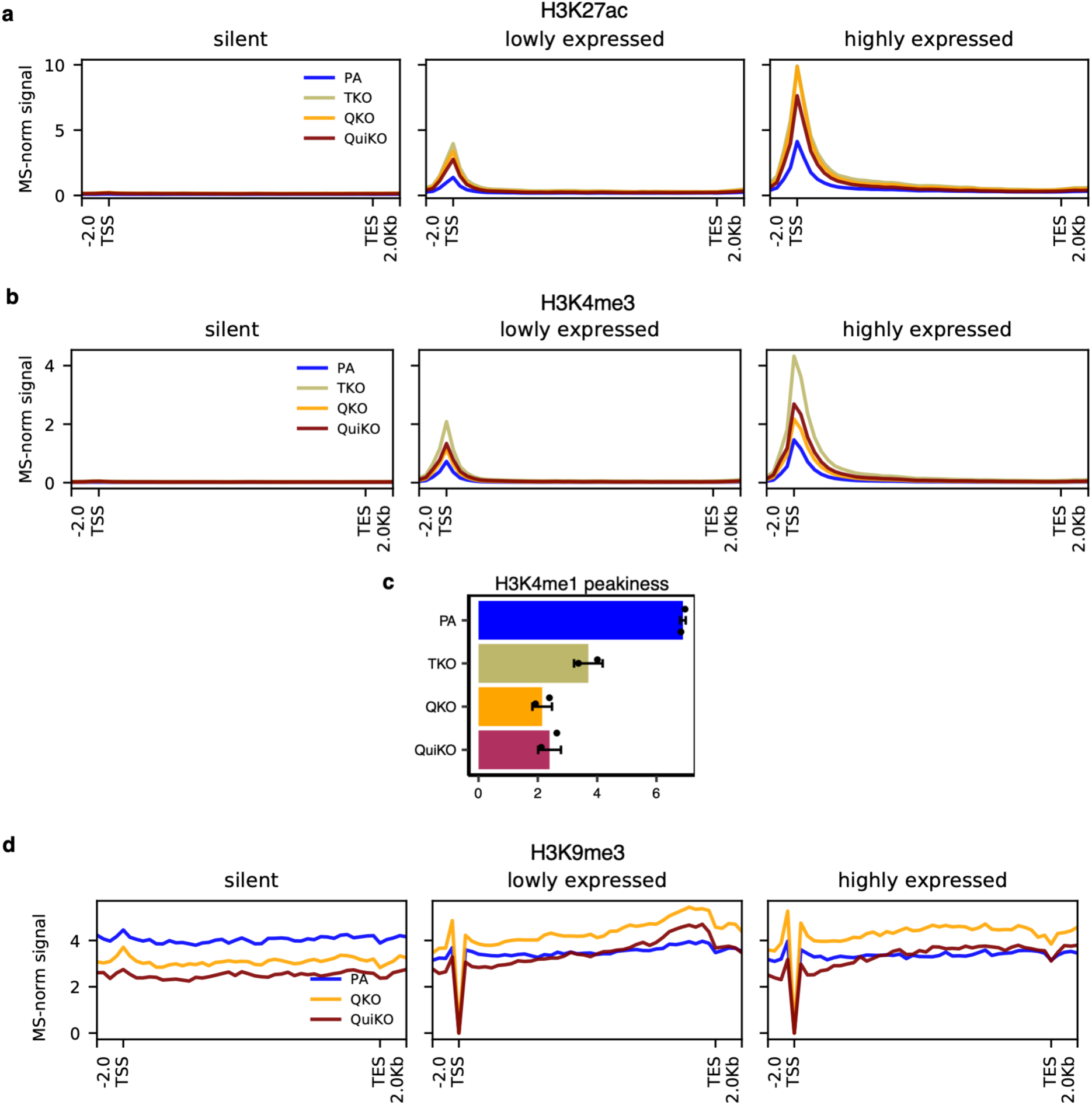
Analysis of other histone marks within gene bodies. **a-b** Gene-body metaplots of MS-normalized H3K27ac and H3K4me3 ChIP-seq signals respectively, illustrating lack of changes within genes (from the same groups of genes in Fig. 3). **c** Barplot of H3K4me1 peakiness scores, showing decreased peakiness in the multi-KOs, which is indicative of less focused H3K4me1 distributions. error bars display the standard deviation around the mean. (n=2 per condition) **d** Gene-body metaplots of MS-normalized H3K9me3 CUT & RUN signal, showing increased H3K9me3 within the gene bodies of lowly and highly expressed genes for QKO compared to parental (PA) cells.

Within the gene bodies of the K36MT mutants, we detected no discernible changes to H3K27ac or H3K4me3, regardless of transcription levels (Supplementary Fig. 4a/b). In contrast, genic H3K4me1 increases substantially, where we observe that cells with greater H3K36me2 depletion experience a greater invasion of H3K4me1, with little dependence on transcription, including in previously silent genes (Fig. 4b). Similarly, we observe by MS that the global abundance of H3K4me1 increases in the multiple-KO conditions (Fig. 4c). Furthermore, the distribution of H3K4me1 becomes increasingly diffuse across the multiple-KO conditions, as indicated by decreasing “peakiness” scores (Supplementary Fig. 4c). This prompted us to investigate potential changes to other histone marks.

Surprisingly, we found significant changes to the genic levels of the silencing-associated modification H3K9me3, whose relationship with H3K36me is currently unclear. In parental cells, H3K9me3 is generally excluded from transcribed genes, however, gradual removal of H3K36me results in an increase of H3K9me3 that appears to be correlated with both gene expression and the amount of residual H3K36me2/3 (Fig. 4d, Supplementary Fig. 3a). In DKO cells, where the loss of NSD1 and NSD2 substantially reduces H3K36me2, genic H3K9me3 increases, especially in lowly expressed genes (Fig. 4d; second box). In TKO cells, which also lack SETD2 and thereby most H3K36me3, the gene body levels of H3K9me3 increase further and appear to lose their dependence on transcription (Fig. 4d). These results intriguingly suggest that the presence of H3K36me2/3 within genes may antagonize the deposition of H3K9me3 in these regions. Given the considerable increase of genic H3K9me3 in the TKO cells, we used MS to quantify the global abundance of H3K9me3 across conditions. Interestingly, we find that the bulk levels of H3K9me3 increase following the depletion of H3K36me3 in both SETD2-KO and TKO cells (Fig. 4e), suggesting that H3K36me3 in particular may antagonize the deposition of H3K9me3 within genes. However, the genome-wide abundance of H3K9me3 in the QKO and QuiKO conditions return to levels comparable to parental cells, and do not necessarily reflect the observed increases in active genes (Fig. 4e, Supplementary Fig. 4d). Overall, despite the increase of H3K9me3 in actively-transcribed genes, the genome-wide abundance in the multiple-KO conditions does not significantly change. Thus, we hypothesized that there must be a corresponding decrease in other regions of the genome.

### Absence of H3K36me leads to redistribution of H3K9me3 from heterochromatin to euchromatin

In parental mMSCs, the distribution of H3K9me3 follows two characteristic patterns: at a very broad scale, H3K9me3 is deposited in large domains reaching several megabases in size (Fig. 5a). These correspond to highly heterochromatic, transcriptionally silent, lamina-associated domains^6,41,42^. At a finer scale, peaks of H3K9me3 are present at the promoters of some silent genes (Fig. 4d; see specifically ‘silent genes’) and transposable elements (TEs). Remarkably, in the H3K36me deficient conditions, the broad H3K9me3 domains entirely disappear (Fig. 5a). The depletion of heterochromatic domains is accompanied by an increase of H3K9me3 in predominantly active regions, which upon closer examination appear to be mostly gene bodies and TEs (Fig. 4d, Supplementary Fig. 5a). To identify and further characterize the genomic compartments that exhibit the highest loss of H3K9me3, we subdivided the genome into 100-kb bins and compared parental cells to the TKO condition, which is largely devoid of H3K36me2/3. The H3K9me3 profiles of the genomic bins subdivide into two distinct clusters: cluster A (orange) corresponds to regions that gain H3K9me3 in TKO cells, which are predominantly genic, while cluster B (blue) represents predominantly intergenic regions that lose H3K9me3 in TKO cells (Fig. 5b). To better represent the large megabase-sized domains of H3K9me3, 100-kb bins within 1-mb were merged together for each cluster. Afterwards, regions in cluster B overlapping with regions in cluster A were excluded. In support of our previous observations, we find that there is a significant gain of H3K9me3 in cluster A regions and a significant loss in cluster B regions (Fig. 5b, c).

**Fig. 5.**
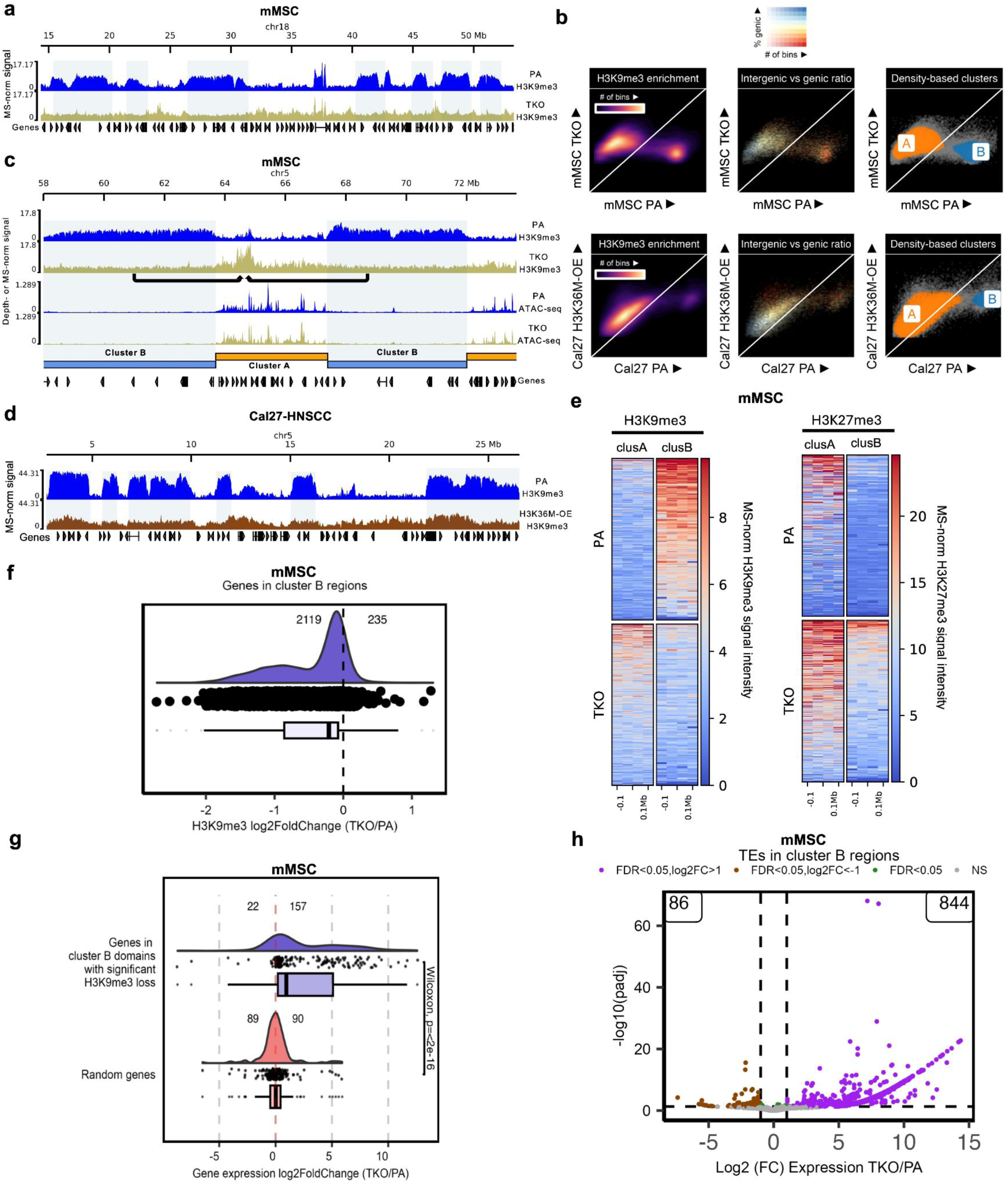
Loss of H3K36me leads to a redistribution of H3K9me3 from heterochromatic regions to euchromatic regions. **a** Genome browser tracks illustrating megabase-sized regions of H3K9me3 loss in 10T-mMSCs TKO compared to PA. **b** Scatterplots of genome-wide H3K9me3 (100-kb resolution) comparing a representative parental sample for mMSCs and HNSCC-Cal27 cell lines against TKO and H3K36M-OE samples respectively, indicating the formations of two clusters: cluster A in which regions gain H3K9me3 and cluster B in which regions lose H3K9me3. **c** Genome browser tracks indicating that in heterochromatic regions, H3K9me3 is lost (designated as cluster B regions), while in euchromatic regions, H3K9me3 is gained (designated as cluster A regions). **d** Genome browser tracks illustrating a similar loss of large H3K9me3 domains in Cal27 OE-H3K36M. **e** Heatmaps centered on cluster A and cluster B regions, illustrating loss of H3K9me3 in cluster B regions, gain of H3K9me3 in cluster A regions, and spreading of H3K27me3 in both cluster A and B regions in the TKO mMSCs. **f** Log2 fold changes of H3K9me3 signal comparing TKO to PA in mMSC, showing that over 90% of genes in cluster B regions (2119/2354) lose H3K9me3 in their gene bodies. **g** Log2 fold changes of gene expression comparing TKO to PA in mMSC, showing that most of the genes in cluster B regions with significant H3K9me3 loss in their gene bodies become upregulated (157 upregulated versus 22 downregulated) whereas a randomized, control set of genes have similar number of genes up- and downregulated (90 and 89 respectively). Statistical significance comparing the two sets of genes was derived from Wilcoxson signed rank test. **h** Volcano plot showing a large number of transposable elements become upregulated in regions of H3K9me3 loss (cluster B domains) in the TKO mMSCs. In the boxplots in **f** and **g**, boxes span the lower (first quartile) and upper quartiles (third quartile), median is indicated with a center line and whiskers extend to a maximum of 1.5 times the interquartile range. Normalized signals were either depth-normalized (ATAC-seq) or MS-normalized (all other tracks), as previously described.

**Supplementary Figure 5.**
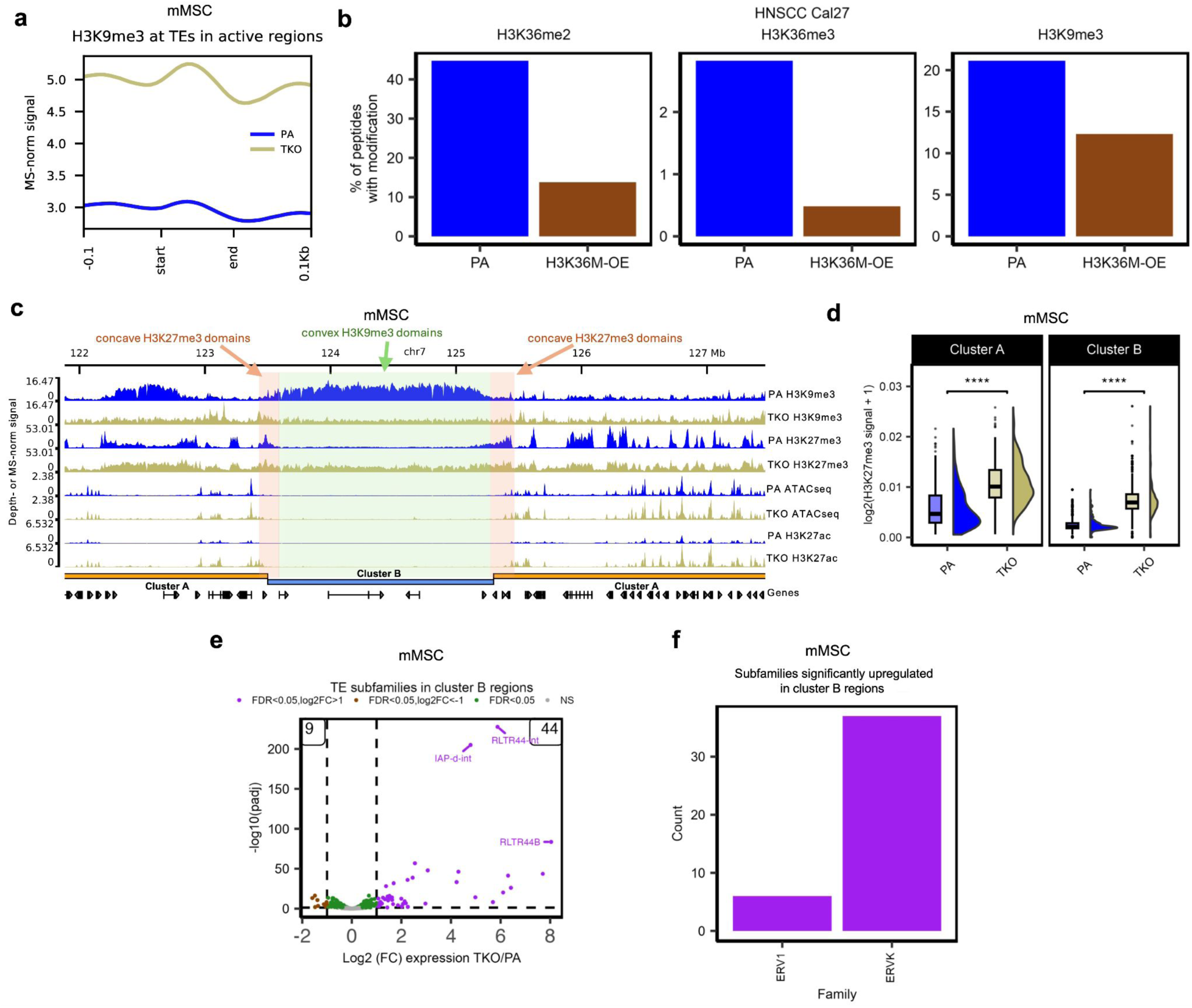
Analysis of H3K9me3 re-distribution and downstream effects following H3K36me depletion. **a** Aggregate plot of MS-normalized H3K9me3 ChIP-seq centered on transposable elements (TEs), showing an increase of H3K9me3 in TEs at active genomic regions. **b** Bar plots of genome-wide prevalence of modifications based on MS, illustrating a global decrease in abundance of H3K36me2/3 and H3K9me3 in the H3K36M-OE cells compared to the PA/wildtype cells in the HNSCC Cal27 cell lines. **c** Representative genome-browser snapshot illustrating that in the absence of H3K36 methylation in the TKO cell line, H3K9me3 is lost and those domains become populated by H3K27me3. Normalized signals were either depth-normalized (ATAC-seq) or MS-normalized (all other tracks), as previously described. **d** Box- and violin-plots of ChIP-seq H3K27me3 signal at cluster A and cluster B regions, indicating a significant increase of H3K27me3 in both regions comparing TKO to PA in mMSCs. **** refers to *p*-value <= 0.0001 from Wilcoxon signed-rank test. **e** Volcano plot of TE subfamilies in cluster B regions, illustrating many more TE subfamilies become upregulated than downregulated in the TKO mMSC. **f** Barplot of subfamilies significantly upregulated in cluster B regions in TKO mMSC, indicating these upregulated TEs largely derive from the ERV1 and ERVK families of TEs.

To verify whether this previously unobserved relationship between H3K36 and H3K9 methylation exists in other cell types, we interrogated the head and neck squamous cell carcinoma (HNSCC) cell line Cal27. Cal27 cells belong to the HPV(-) subgroup of HNSCC but harbor no endogenous mutations affecting H3K36me^18^. In these cells we overexpressed the H3K36M oncohistone, a mutation that globally reduces all H3K36me states and that is found in the HPV(-) HNSCC patient cohort and in other cancers (Supplementary Fig. 5b)^14,21,43^. Similar to mMSCs, we find that Cal27 parental cells contain well-defined H3K9me3-marked heterochromatic domains (Fig. 5d). Strikingly, upon overexpressing the H3K36M mutation, we observe a drastic depletion of the large H3K9me3 domains (Fig. 5d), corresponding to cluster B regions in our genome-wide density-based cluster analysis of the Cal27 cell lines (Fig. 5b; 2nd row). Interestingly, unlike in mMSCs, we observe a nearly two-fold depletion of H3K9me3 by MS in the Cal27 H3K36M-OE cells (Supplementary Fig. 5b).

In parental mMSCs, large heterochromatic, gene-poor regions are characterized by “convex” H3K9me3 domains, flanked by reciprocal “concave” domains of H3K27me3 (Supplementary Fig. 5c). These regions are transcriptionally inactive with no open chromatin sites or other active epigenetic modifications (Supplementary Fig. 5c). In the absence of H3K36 methylation in the TKO cell line, H3K9me3 is lost and those domains become populated by H3K27me3 (Supplementary Fig. 5c; Fig. 5e: 2nd heatmap), possibly as a compensatory silencing mechanism. Overall, in addition to invading gene bodies found in active, gene-rich regions (cluster A), H3K27me3 also spreads into inactive, gene-poor regions where H3K9me3 is lost (cluster B), suggesting a shift from a constitutive to a facultative heterochromatic state in these regions (Fig. 5e; Supplementary Fig. 5d).

Nevertheless, the silencing compensation by H3K27 methylation appears to be inadequate. In cluster B domains, approximately 90% (2119/2354) of genes lose H3K9me3 (Fig. 5f). Of these 2119 genes, those with the most significant loss of H3K9me3 become upregulated (157/179 genes) (Fig. 5g). Furthermore, we find that many of the TEs in cluster B regions become differentially expressed, with the majority becoming upregulated (844 up versus 86 down) (Fig. 5h). Further characterization of these TEs reveals that 44 specific sub-families become upregulated in comparison to 9 that become downregulated (Supplementary Fig. 5e), with most of the upregulated TEs belonging to the ERV family (Supplementary Fig. 5f). Our findings indicate that both genes and TEs in cluster B regions become de-repressed following the loss of H3K9me3, suggesting an overall de-repression of heterochromatin.

At this time, we can only speculate regarding the mechanisms of the redistribution of H3K9me3 following the depletion of H3K36me. It may be a primary effect, such as diffusion of H3K9 methyltransferases (e.g. SUV39H1/2) away from heterochromatin and into active genic regions, or a secondary effect, such as general epigenome decompartmentalization. However, we expect that the loss of large H3K9me3 marked domains leads to additional epigenetic changes in these heterochromatic regions.

### Downstream epigenetic cascade following the depletion of H3K9me3 decorated heterochromatin

We further investigated the epigenetic changes that occur, in the absence of H3K36me, within the heterochromatic regions now devoid of large H3K9me3 domains. Strikingly, we observe changes in chromatin accessibility, as indicated by ATAC-seq and the presence of new peaks within previously inaccessible chromatin (1914 new open chromatin regions in the TKO cells compared to parental) (Fig. 6a, b). Consistent with H3K27me3 being associated with facultative heterochromatin, this suggests that the gain of H3K27me3 within cluster B regions is insufficient to maintain a constitutive heterochromatic state (Fig. 6a, Supplementary Fig. 5c). Interestingly, most of these open chromatin peaks are very small in comparison to those typically found in transcriptionally active regions, such as active promoters and enhancers. The gain of open chromatin is accompanied by a significant increase of H3K4me1, and in some regions, H3K27ac (Fig. 6c). This further explains the increase in the overall abundance of H3K4me1 observed by MS, which in addition to increasing within the gene bodies of H3K36me deficient cells (Fig. 4c), also appears to increase in regions that lose H3K9me3 (Fig. 6c). Furthermore, we observe a decrease of DNAme at these sites, indicating a release from silencing (Supplementary Fig. 6a). Lastly, by RNA-seq, we observe an increase in transcription within newly open chromatin, further supporting that these regions are no longer constitutively repressed (Fig. 6c; fourth heatmap). Subsequent analyses of genes previously occupied by H3K9me3 reveals new ATAC-seq peaks at their promoters (Fig. 6a; see gene *Sprr1a* as an example) and a dramatic increase of expression in comparison to a control set of genes with comparable expression levels (Supplementary Fig. 6b, c).

**Fig. 6.**
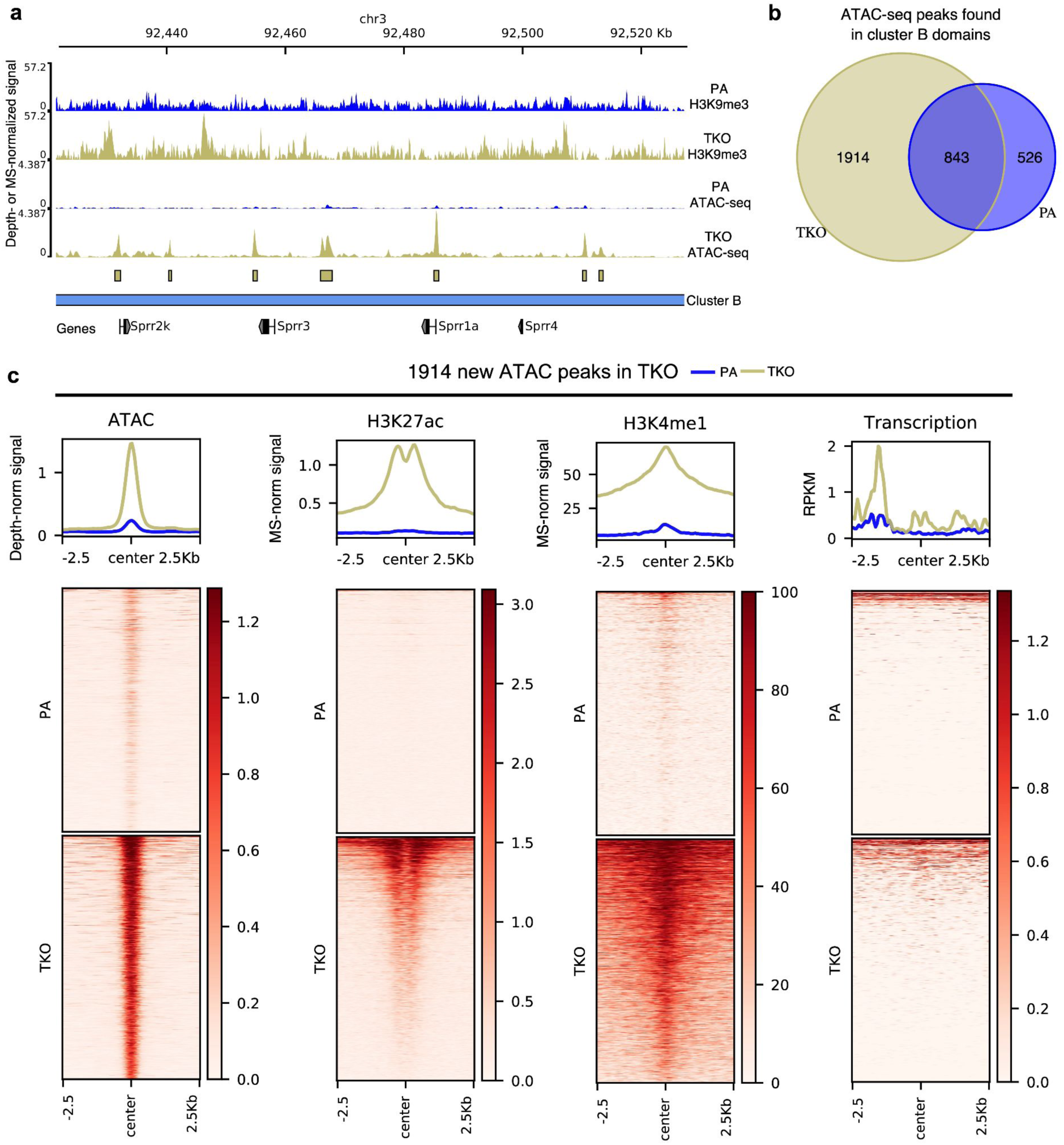
Presence of new open chromatin peaks within regions of H3K9me3 loss. **a** Genome browser tracks depicting opening of new ATAC-seq peaks within cluster B regions, where H3K9me3 is lost in TKO. **b** Venn diagram of ATAC-seq peaks found in cluster B regions, where 1914 new ATAC-seq peaks are found exclusively in TKO. **c** Heatmaps centered on the 1914 new ATAC-seq peaks opening in TKO, indicating an upregulation of active marks (H3K27ac and H3K4me1) and gene expression activity. Normalized signals were either depth-normalized (ATAC-seq), MS-normalized (for H3K27ac and H3K4me1) or RPKM (reads per kilobase per million mapped) transformed.

**Supplementary Figure 6.**
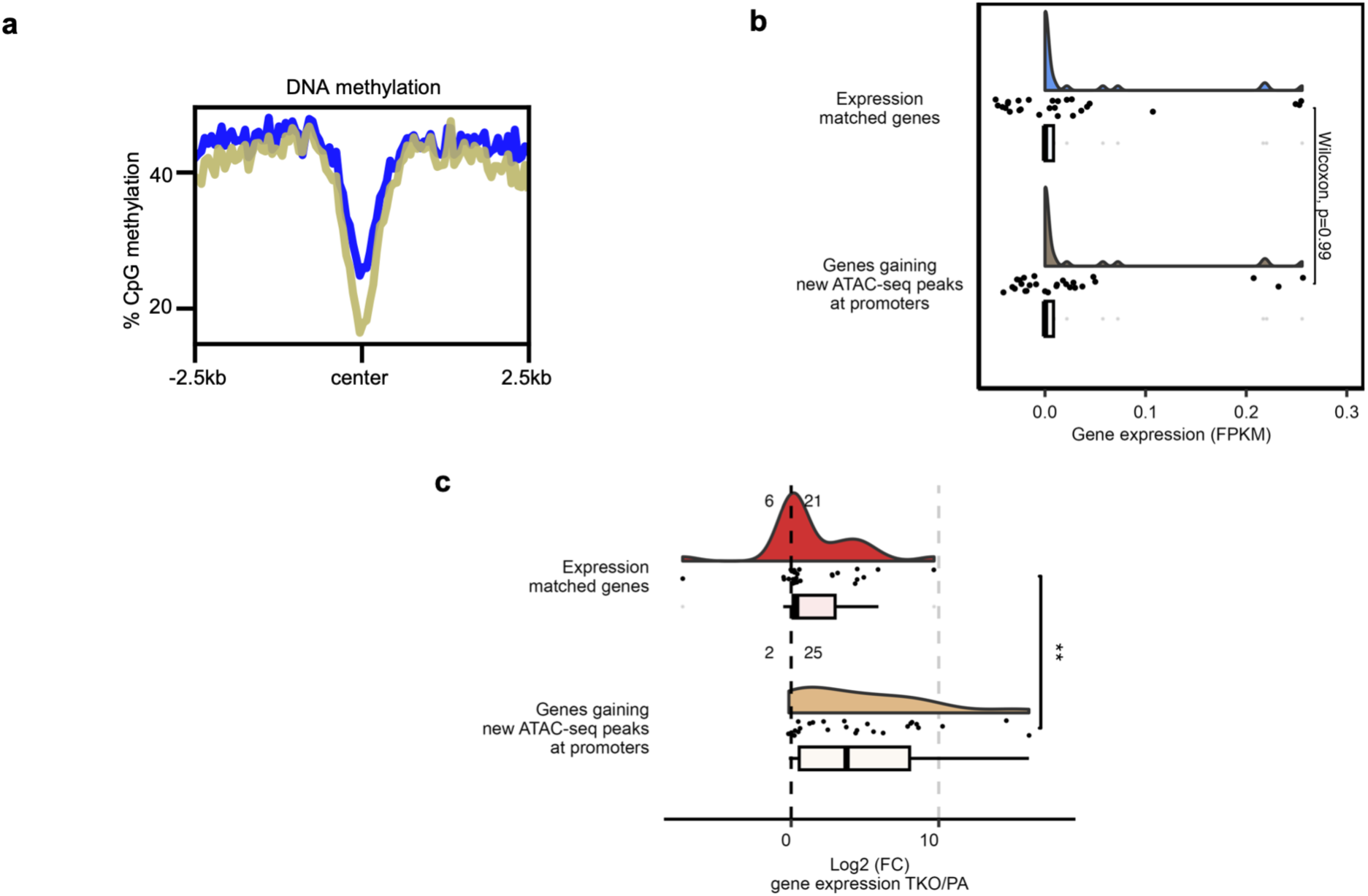
Changes in DNA methylation and gene expression in cluster B regions with newly open chromatin in TKO mMSCs. **a** Aggregate plot of DNA methylation at CpG sites centered on the 1914 peaks opening in TKO, indicating a downregulation of CpG methylation at these new ATAC-seq peaks. **b** Gene expression plots showing no significant difference in basal expression (FPKM) for genes gaining new ATAC-seq peaks at their promoters in the TKO cells compared to a control set of genes with similar gene expression. Basal expression refers to gene expression in the PA/wildtype cells. **c.** Gene expression plots showing that 23/25 genes that gained new ATAC-seq peaks at their promoters in the TKO cells had upregulated gene expression compared to a control set of expression-matched genes from **b. \*\*** refers to p-value <= 0.01 from a Wilcoxon signed-rank test. For **b** and **c,** boxes span the lower (first quartile) and upper quartiles (third quartile), median is indicated with a center line and whiskers extend to a maximum of 1.5 times the interquartile range.

### Absence of H3K36me results in a drastic reduction of nuclear compartmentalization

At a basic level, chromatin can be segregated into active and inactive compartments, which previous studies have classified as A and B compartments, respectively^44^. The severe changes to both euchromatic and heterochromatic histone modifications resulting from the depletion of H3K36me prompted us to investigate possible changes to chromatin conformation and nuclear compartmentalization. To do this, we performed Hi-C on parental and QuiKO mMSCs, which lack all H3K36me. The parental mMSCs have distinct A/B compartments and euchromatin/heterochromatin structure, as illustrated by the strong checkerboard pattern at the chromosomal scale (Fig. 7a; upper diagonal). The inactive B compartments are highly correlated with H3K9me3 domains (Supplementary Fig. 7a) and show significant B-B compartment interactions (Fig. 7b), which may be interpreted as strong heterochromatin compaction. Notably, the B-B compartment interactions are nearly twice as frequent as A-A interactions (Fig. 7b). In the absence of H3K36me (QuiKO) and following the loss of H3K9me3 domains, we observe a drastic loss of genome-wide compartmentalization (Fig. 7a; lower diagonal). This is manifested by a 2-fold reduction in B-B and A-A compartment interactions (Fig. 7b) and a homogenization of compartment scores resulting in the loss of the bimodal nature of compartment identity, especially for B-compartments (Fig. 7c). The overall number of compartments is comparable between conditions, although there is a slight shift from A to B compartments (Fig. 7d). Nevertheless, further investigation into the global loss of these heterochromatic interactions reveals that long-range interactions (> 10 mb) are particularly diminished within chromosomes (*cis*-interactions) (Supplementary Fig. 7b). We also assessed changes to topologically associating domains (TADs) and chromatin loops. Interestingly, we found no significant changes to the total number of TADs nor the strength of TAD boundaries (Supplementary Fig. 7c, d); however, in QuiKO cells we do observe over a two-fold loss in the total number of loops in comparison to parental cells (1448 in parental compared to 701 in QuiKO) (Supplementary Fig. 7e). Furthermore, we observe that a greater proportion of long-range loops (loops spanning distances greater than 500 kb) are diminished compared to short-range loops (Supplementary Fig. 7f).

**Fig. 7.**
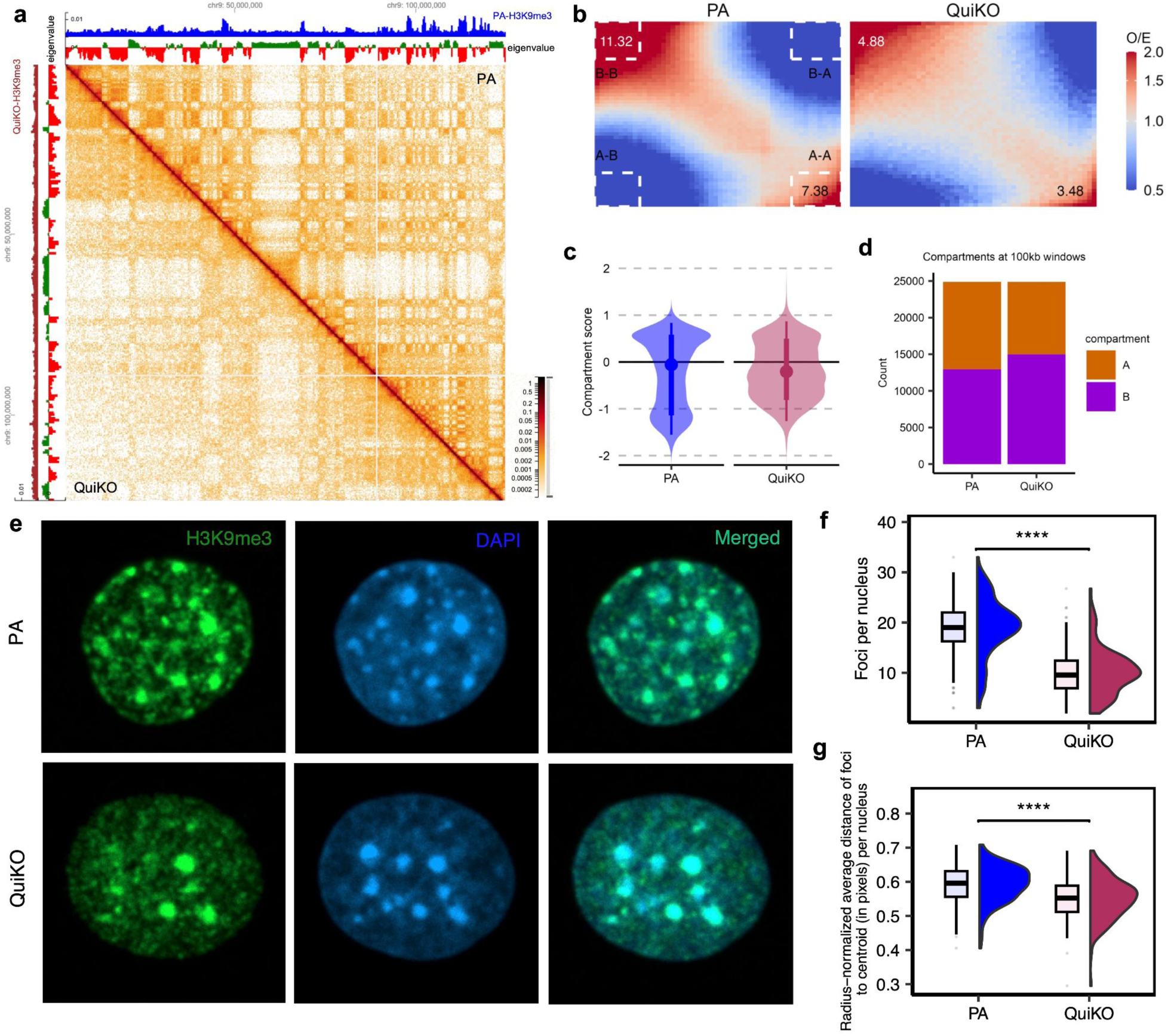
Depletion of H3K36me results in decreased nuclear compartmentalization. **a** Representative Hi-C heatmaps for parental (PA) (top right of diagonal) and QuiKO (bottom left of diagonal) cells, illustrating a reduction of nuclear compartmentalization following complete depletion of H3K36me. The blue and maroon tracks correspond to MS-normalized H3K9me3 for PA and QuiKO cells respectively. The 1st eigenvalue is shown, with positive and negative eigenvalues values corresponding to active A (in green) and inactive B compartments (in red) respectively. **b** Saddle plots showing decreased A-A and B-B *cis* compartment interactions in QuiKO compared to PA. O/E refers to observed over expected scores. A-A interactions are in the lower right corner whereas B-B interactions are in the upper left corner. **c** Violin plot showing a shift of compartment scores (i.e. the first eigenvalues as calculated in *cis*-chromosomal data) towards 0, indicating a decrease in compartmentalization in QuiKO cells compared to PA. **d** Bar plot illustrating a shift of A to B compartments in QuiKO. **e** Representative immunofluorescence images, depicting a reduction of H3K9me3 foci, especially those at the periphery, in QuiKO compared to PA nuclei. Nuclei were stained with anti-H3K9me3 antibody and DAPI respectively. For proper comparison of the quantity and spatial distribution of the foci, all images were taken under the same conditions and similar fluorescence intensity was ensured in each image. **f** Boxplots showing a reduction of foci per nuclei in QuiKO cells compared to PA. The foci count per nucleus in the QuiKO cells were normalized by the ratio of the mean nuclei surface area of the PA cells (13747.99 pixels) to that of the QuiKO cells (14394.36 pixels). **g** Boxplots of the average distance of foci to the center of each nucleus, showing that the foci in the PA cells are generally further away from the center than in the QuiKO cells. The average distance of the foci per nucleus was normalized by the radius of each nucleus. For **f** and **g,** n=130 nuclei for each condition and **** represents p-value <= 0.0001 from a Wilcoxon signed-rank test.

**Supplementary Figure 7.**
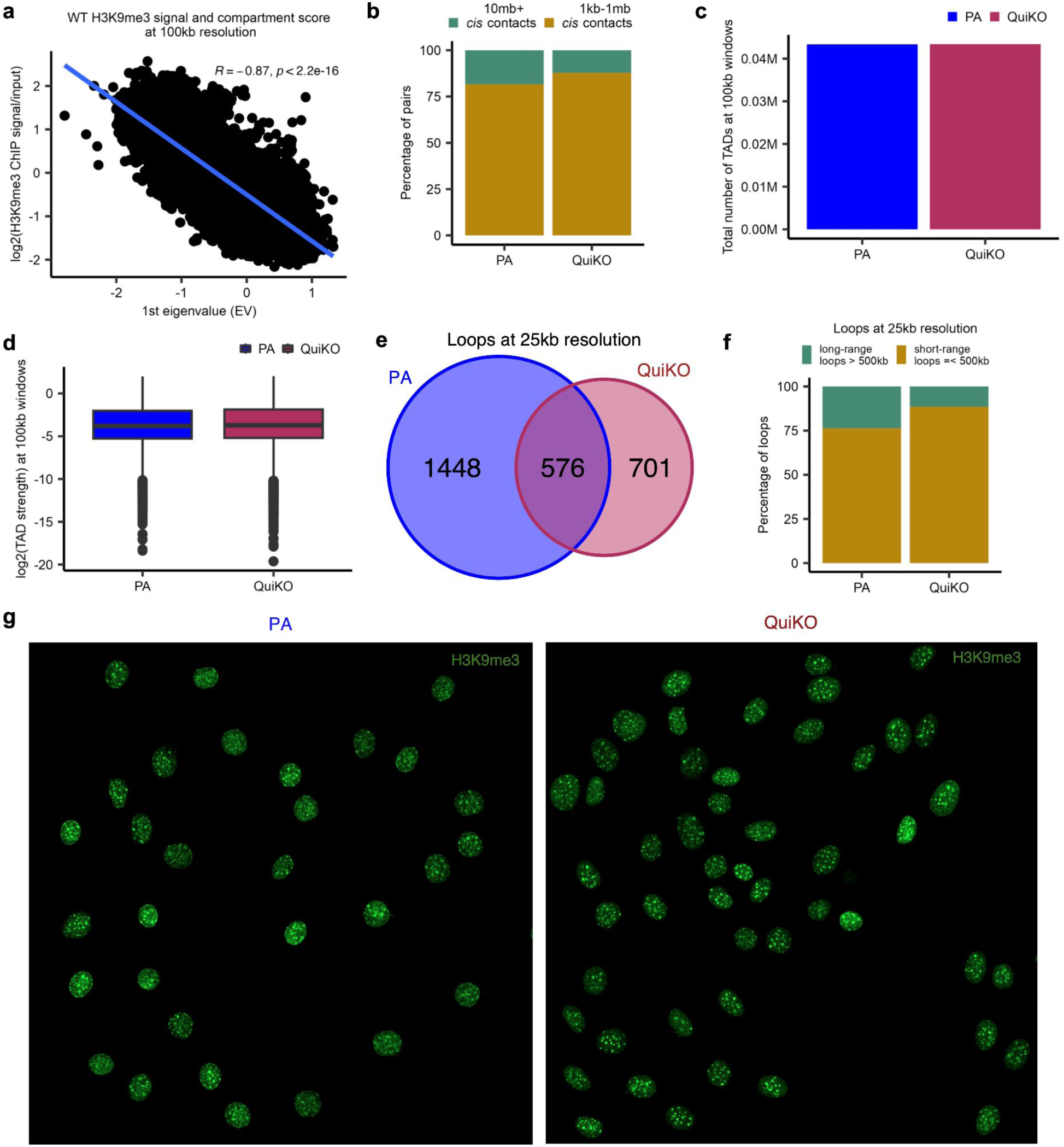
Analysis of chromatin structure following total H3K36me2 depletion. **a** Correlation plot depicting a strong negative linear correlation between H3K9me3 ChIP signal and the 1st eigenvalue (compartment score). Reported values are Pearson’s correlation coefficient (*R*) and the associated *p*-value. **b** Barplots of long-range *cis*-contacts, showing a reduction of very long-range contacts (10 mb+) in the QuiKO cells compared to the PA cells. **c** Barplots depicting the total number of TADs at 100kb windows, with no significant differences between the PA and QuiKO cells. **d** Boxplots of TAD strengths at 100kb windows, showing no significant differences between PA and QuiKO cells. **e** Venn diagram of the number of loops at 25 kb resolution, depicting decreased number of loops in the QuiKO cells compared to the PA cells. **f** Barplots showing a greater proportion of long-range loops (loops spanning distances greater than 500 kb) are diminished compared to short-range loops in QuiKO compared to PA cells. **g** Representative immunofluorescence images depicting loss of H3K9me3 foci at the periphery, with the remaining foci clustered towards the center in QuiKO nuclei compared to PA nuclei. Nuclei were stained with anti-H3K9me3 antibody. For proper comparison of the quantity and spatial distribution of the foci, all images were taken under the same conditions and similar fluorescence intensity was ensured in each image.

Our analyses suggest that the most prominent changes are the dissolution of compartment interactions, especially long-range interactions, and the loss of heterochromatin compaction. To physically observe these alterations at the microscopic level, we performed immunofluorescence (IF) to visualize the distribution of H3K9me3 (Supplementary Fig. 7g). In the parental cells, H3K9me3 forms numerous distinct foci, which likely correspond to large heterochromatic domains (Fig. 7e, f; Supplementary Fig. 7g). Many of those foci locate to the nuclear periphery (Fig. 7e), suggesting interaction with the nuclear lamina that is characteristic of heterochromatin^45,46^. In the QuiKO cells, H3K9me3 shows a more diffused distribution with fewer, distinct foci (Fig. 7e, f) and a much weaker localization at the nuclear envelope, with the remaining H3K9me3 foci located more towards the centre of the nucleus (Fig. 7e, g; Supplementary Fig. 7g). Under both conditions, H3K9me3 co-localizes with DAPI, which is known to preferentially stain AT-rich regions (Fig. 7e). However, in QuiKO cells, the DAPI signal also shifts towards the centre of the nucleus, further supporting significant changes to chromatin organization (Fig. 7e). Overall, our results demonstrate that the depletion of H3K36me affects not only euchromatin, but results in massive overall decompartmentalization, reduced long range heterochromatic interactions, and a shift of H3K9me3 signal away from the nuclear envelope and towards the nuclear centre.

## Discussion

H3K36 methylation is emerging as a key factor shaping the euchromatic epigenome^2,14,35,47–50^. While the genomic patterns of H3K36me deposition have already been characterized^2,18,22^, its function remains far from being understood. One of the difficulties lies in the broad nature of the distribution: H3K36me2 is present in most euchromatic intergenic regions and transcribed genes while H3K36me3 is present in transcribed gene bodies, where it usually co-localizes with H3K36me1/2. Our system, consisting of combinatorial and stepwise deletion of the five major K36MTs provides a unique opportunity to gradually deplete H3K36me from different genomic compartments and observe the downstream effects of H3K36me removal on the epigenome and transcriptome. In particular, the TKO, QKO, and QuiKO conditions allow us to investigate the role of H3K36me2 in specific and localized settings. TKO cells exhibit H3K36me2 in relatively narrow peaks surrounding active promoters and enhancers, and we have previously found that NSD3 is responsible for their deposition. QKO cells retain only a handful of very specific peaks at the CREs of a small subset of developmentally important genes, and QuiKO cells have no detectable H3K36me. Therefore, one of our primary goals was to define the role of H3K36me2 in active regulatory regions.

Interestingly, we find that the depletion of H3K36me2 has a more profound impact on the activity of enhancers in comparison to promoters, as indicated by the greater reduction of activating histone marks and chromatin accessibility at enhancers. Furthermore, we find that enhancers, compared to promoters, experience greater encroachment of the silencing modification, H3K27me3. Accordingly, we find that the majority of genes whose expression relies on H3K36me2-marked enhancers decrease in expression upon the loss of H3K36me2. Overall, the observed changes to gene expression can be broken down into two parts: H3K36me2-dependent changes, and secondary, downstream effects from the global depletion of H3K36me2 and/or ablation of the respective K36MTs. The latter, in part, has been documented in studies characterizing non-enzymatic properties of the K36MTs, and this is further supported by the existence of isoforms lacking catalytic activity, such as NSD2-short and NSD3-short ^23,48,51^. Furthermore, we observe that in the individual NSD3- and ASH1L-KO cells, the levels of H3K36me2 remain largely unchanged at the H3K36me2-dependent target genes identified in TKO and QKO cells, likely resulting from compensation and/or partial redundancies by NSD1 or NSD2, which appear to have less stringent dependencies on the localization of their catalytic activities. When considering only genes whose enhancers were marked by H3K36me2, our results suggest that the decrease in gene expression following the loss of H3K36me2 can largely be attributed to reduced enhancer activity, as indicated by the greater reduction of activating histone marks and greater increase of H3K27me3. These results support previous studies in other systems: in mouse embryonic stem cells, NSD1 was found to be more highly enriched at enhancers in comparison to promoters, while in patient HNSCC cells, the depletion of H3K36me2 resulting from NSD1 loss of function mutations was found to have the greatest effect on reducing the enhancer activity of affected target genes^2,18,23^. Collectively, our results support that the presence of H3K36me2 at CREs predominantly influences transcription via its effect on enhancers. We next focused on the effect of H3K36me depletion in gene bodies. Expressed genes normally maintain an open chromatin structure, which is mostly devoid of silencing-associated modifications, such as H3K27me3 and H3K9me3. It is generally believed that the presence of H3K36me3 within expressed genes excludes the deposition of H3K27me3^30,31^. There is strong evidence that H3K36me3 prevents binding of EZH2 to the already modified nucleosome, and the two marks are mutually exclusive on the same histone tail. H3K36me2 has a similar, albeit less pronounced inhibitory effect on the deposition of H3K27me3. We are not aware of any previous studies that have specifically examined the effect of H3K36me on the genic exclusion of H3K27me, *in vivo*. Starting with H3K36me3, we find that although its presence does result in the strongest exclusion of H3K27me2/3 from expressed genes, in SETD2-KO cells, which lack nearly all H3K36me3, we find only a modest increase of genic H3K27me3. Subsequent ablation of the NSD proteins further demonstrates that H3K36me2 provides an additional level of protection against H3K27me3. However, even in the total absence of H3K36me, transcribed gene bodies do not become fully invaded by H3K27me2/3. We propose that the remaining barrier is the result of histone turnover during transcription and the resulting dilution of H3K27me-marked nucleosomes, and possibly the presence of DNAme, which is known to prevent binding of PRC2. Interestingly, the total absence of H3K36me in the mMSCs does not result in a marked reduction of DNAme within gene bodies, showing that, in this cell type and possibly other differentiated cells, the activity of maintenance DNMT1 is sufficient to maintain DNA methylation, even in the absence of proper targeting of *de novo* DNMT methyltransferases (Fig. 4a).

Unexpectedly, we find that cells lacking H3K36me show a significant invasion of the other major silencing modification, H3K9me3, into transcribed gene bodies. This is surprising, since H3K36me has no known relationship with H3K9me3. A putative co-existence of H3K36me3 and H3K9me3 has previously been reported, but only in the context of specific promoter and enhancer occupancy^52^. It is unclear how, mechanistically, the invasion of H3K9me3 into transcribed genes is prevented, but we found it to be highly associated with the depletion of H3K36me2 and H3K36me3. In the absence of H3K36me, both highly transcribed genes and intergenic regions acquire nearly uniform levels of H3K9me3. The partial invasion of both H3K27me3 and H3K9me3 into gene bodies results in a significant reduction of gene expression, at least in genes with the greatest invasion of these suppressive marks.

Lastly, since the overall abundance of H3K9me3 appears nearly unchanged in the multiple-KO conditions, the genic increase must be the result of a redistribution of this modification from other regions of the genome. Accordingly, we found that the genic increase is accompanied by a massive loss of large intergenic heterochromatic H3K9me3 domains. To demonstrate that this trend is not limited to a single cell type, we overexpressed the H3K36M mutant histone in the HNSCC cell line Cal27, and similarly found that the reduction of genome-wide H3K36me was accompanied by the loss of large H3K9me3 domains and a gain of isolated peak signal and genic levels. This relationship between H3K36me and H3K9me3 is a novel finding, which appears to be generalizable and merits further investigation.

The disappearance of the broad heterochromatic H3K9me3 domains is accompanied by a possible compensatory replacement with H3K27me3 (Fig. 5e, Supplementary Fig. 5c). However, it appears that the compensation is not complete, since we observe the appearance of new open chromatin peaks, accompanied by an increase of H3K27ac and H3K4me1, indicating that the previously inaccessible and silenced regions gain bivalent and/or active characteristics, further accompanied by some slight de-repression of previously silenced genes and regulatory elements.

Finally, following the complete loss of H3K36me in QuiKO cells, we observe a drastic loss of nuclear compartmentalization. Specifically, inactive compartments appear to lose their structural integrity, consistent with the loss of broad H3K9me3 domains. Furthermore, there is a reduction in long-range intra-chromosomal contacts and chromatin loops. Collectively, this suggests a crucial role for H3K36me in supporting chromatin architecture, possibly by helping to establish or maintain proper segmentation of genomic compartments. Following the loss of H3K36me, many of these compartment boundaries disappear and a substantial reorganization of the epigenome is observed, resulting in significant transcriptomic changes: H3K9me3 redistributes from the nuclear periphery and into centrally located euchromatic regions (Fig. 7a, e), while previously inaccessible regions gain characteristics associated with active compartments (Fig. 6c). Previous studies by our group and others have identified enrichment of H3K36me at insulator elements, notably CTCF; thus, assessing whether changes occur to such insulators in the absence of H3K36me may provide additional mechanistic insights^18,22,29^.

H3K36me continues to emerge as an essential component of chromatin regulation throughout development, and pathological mutations affecting this regulation are frequently implicated in developmental disorders and cancer. Its presence influences not only the deposition of other epigenetic modifications and gene expression, but our study demonstrates that this influence extends to the regulation of genomic organization and higher order chromatin structure. Our findings uncover many intriguing avenues that merit further investigation. The relationship between H3K36me, H3K9me3 and chromatin architecture is novel and may provide valuable therapeutic and/or mechanistic insights into the numerous developmental disorders and cancers affected by the aberrant regulation of H3K36me. Since both H3K36me2/3 appear to contribute to the observed changes, it will be paramount to dissect the individual contributions of the K36MTs to these phenomena.

## Materials and methods

### Cell culture

C3H10T1/2 mouse mesenchymal stem cells (ATCC) were cultured in DMEM (1X) (Invitrogen) with 10% fetal bovine serum (FBS) (Wisent) and supplemented with 1% GlutaMax. C3H10T1/2 cell lines harboring individual knockouts of NSD1, NSD2, NSD3, SETD2, and ASH1L, as well as the NSD1/2/3-SETD2-QKO and NSD1/2/3-SETD2-ASH1L-QuiKO cell lines were established by our group as described in^22^. C3H10T1/2 cells overexpressing the H3K36M mutation were provided by the lab of Dr. Chao Lu (Stanford University), and C3H10T1/2 NSD1/2-DKO and C3H10T1/2 NSD1/2-SETD2-TKO cells were provided by the lab of Dr. C. David Allis (Rockefeller University). Cal27 HNSCC cells (ATCC, CRL-2095) were cultured in DMEM (F12) (Invitrogen) with 10% FBS. *Drosophila* S2 cells were cultured in Schneider’s *Drosophila* medium (ThermoFisher) containing 10% FBS. All cell lines tested negative for mycoplasma contamination.

### Generation of Cal27 HNSCC cells overexpressing the H3K36M mutation

Epitope-tagged H3.3 was previously cloned into pCDH-EF1-MCS-Puro lentiviral vector and site-directed mutagenesis techniques were used to generate the H3K36M mutation^14^. To produce lentivirus, 293T cells were transfected with a lentiviral vector containing H3.3K36M (provided by Dr. Chao Lu) and helper plasmids (psPAX2 and pMD2.G). Supernatant containing lentivirus was collected after 48h, filtered and used to infect Cal27 cells. Transduced cells were grown under puromycin selection (1 μg/ml) for 48 h after transduction.

### Crosslinking and ChIP-seq

Approximately 20 million cells per cell line were used. Crosslinking was performed in 150 mm cell culture plates using 1% formaldehyde (Sigma) at room temperature with gentle rocking for 10 minutes. The crosslinking reaction was quenched using 1.25 M Glycine with gentle rocking at room temperature for 5 minutes. Fixed cell preparations were washed with ice-cold PBS, scraped, and then washed twice more with ice-cold PBS. Crosslinked pellets were resuspended in 500 μl cell lysis buffer (5 mM pH 8.5 PIPES, 85 mM KCl, 1% (v/v) IGEPAL CA-630, 50 mM NaF, 1 mM PMSF, 1 mM phenylarsine oxide, 5 mM sodium orthovanadate, EDTA-free protease inhibitor tablet) and incubated for 30 minutes on ice. Samples were centrifuged and pellets were resuspended in 500 μl of nuclei lysis buffer (50 mM pH 8.0 Tris-HCl, 10 mM EDTA, 1% (w/v) SDS, 50 mM NaF, 1 mM PMSF, 1 mM phenylarsine oxide, 5 mM sodium orthovanadate, EDTA-free protease inhibitor tablet) and incubated for 30 minutes on ice. Sonication of lysed nuclei was performed using the BioRuptor UCS-300 at maximum intensity for 75-90 cycles (10s on, 20s breaks). Sonication efficiency to achieve fragments between 150 - 500 bp was evaluated using gel electrophoresis on a reversed-crosslinked and purified aliquot from each sample. After sonication, chromatin was diluted to reduce SDS levels to 0.1% and concentrated using Nanosep 10K OMEGA (Pall) columns. Prior to the chromatin immunoprecipitation (ChIP) reaction, 2% sonicated *Drosophila* S2 cell chromatin was spiked into each sample for quantification of total levels of histone marks. The ChIP reactions were performed using the Diagenode SX-8G IP-Star Compact and Diagenode automated iDeal ChIP-seq Kit for Histones. Dynabeads Protein A (Invitrogen) were washed, then incubated with specific antibodies (Supplementary Table 1), 1.5 million cells of sonicated cell lysate, and protease inhibitors for 10 hrs, followed by a 20 min wash cycle using the provided wash buffers (Diagenode Immunoprecipitation Buffers, iDeal ChIP-seq Kit for Histones). Reverse cross-linking was then performed using 5M NaCl at 65℃ for 4 hrs. ChIP samples were then treated with 2 μl RNase Cocktail at 65℃ for 30 min followed by 2 μl Proteinase K at 65℃ for 30 min. Samples were then purified with QIAGEN MinElute PCR purification kit (QIAGEN) as per the manufacturer’s protocol. In parallel, input samples (chromatin from about 50,000 cells) were reverse crosslinked and DNA was isolated following the same protocol. Library preparation was performed using the Kapa Hyper Prep library preparation reagents following the manufacturer’s protocol (Kapa Hyper Prep Kit, Roche 07962363001). ChIP libraries were sequenced by the McGill University Genome Centre using the Illumina HiSeq 4000 at 50 bp single reads or NovaSeq 6000 at 100 bp single reads.

### Cut & Run

CUT&RUN reactions were performed with the Epicypher CUTANA ChIC/CUT&RUN Kit Version 3 (cat# 14-1048) using 550,000 fresh C3H10T1/2 cultured cells and following the manufacturer’s protocol (Epicypher User Manual Version 3.5) for adherent cells. Prior to performing CUT&RUN, optimization of C3H10T1/2 cell permeabilization was performed following Appendix 1.1 (Epicypher User Manual Version 3.5). Cell permeabilization was found to be most optimal using a final working concentration of 0.01% Digitonin (Promega). Successful binding of cells to Concanavalin A beads was confirmed as recommended and outlined by the manufacturer. Manufacturer supplied H3K4me3 positive control and IgG negative control antibodies were used and the reactions were performed in parallel with processed samples. Processed samples were incubated with specific antibodies (Supplementary Table 1). Manufacturer also supplied *E.coli* spike-in DNA, which was used in each reaction for sequencing normalization. Quantification of purified CUT&RUN DNA was performed using the Qubit fluorometer with the 1X dsDNA HS Assay Kit (Invitrogen). Library preparations were performed using the Epicypher CUTANA CUT&RUN Library Prep Kit Version 1 (cat# 14-1001) following the manufacturer’s protocol. Final CUT&RUN library concentrations were quantified as previously described, and enrichment of mononucleosome-sized fragments was assessed by Bioanalyzer (Agilent) prior to sequencing. CUT&RUN libraries were sequenced at a depth of 15M reads per sample by the McGill University Genome Centre using the Illumina NovaSeq 6000 at 100bp paired-end reads.

### ATAC-seq

ATAC-Seq library preparation was performed according to the Omni-ATAC protocol ^53^. 50,000 C3H10T1/2 cultured cells were resuspended in 1 ml of cold ATAC-seq resuspension buffer (RSB; 10 mM Tris-HCl pH 7.4, 10 mM NaCl, and 3 mM MgCl2 in water). Cells were centrifuged at 500 rcf for 5 min in a pre-chilled (4°C) fixed-angle centrifuge. After centrifugation, supernatant was aspirated and cell pellets were then resuspended in 50 μl of ATAC-seq RSB containing 0.1% IGEPAL, 0.1% Tween-20, and 0.01% digitonin by pipetting up and down three times. This cell lysis reaction was incubated on ice for 3 min. After lysis, 1 ml of ATAC-seq RSB containing 0.1% Tween-20 (without IGEPAL and digitonin) was added, and the tubes were inverted to mix. Nuclei were then centrifuged for 10 min at 500 rcf in a pre-chilled (4°C) fixed-angle centrifuge. Supernatant was removed and nuclei were resuspended in 50 uL transposition mix (2x TD Buffer, 100 nM final transposase, 16.5 uL PBS, 0.5 uL 1% digitonin, 0.5 uL 10% Tween-20, 5 uL H2O) (Illumina Tagment DNA Enzyme and Buffer Small Kit, 20034197). Transposition reactions were incubated at 37 °C for 30 min in a thermomixer with shaking at 1000 rpm. Reactions were cleaned up with DNA Clean and Concentrator 5 columns (Zymo Research). Illumina Nextera DNA Unique Dual Indexes Set C (Illumina, 20027215) were added and amplified (12 cycles) using NEBNext 2x MasterMix. Sequencing of the ATAC-Seq libraries was performed on the Illumina NovaSeq 6000 system using 100-bp paired-end sequencing.

### RNA-seq

Total RNA was extracted from approximately 1 million cells using the AllPrep DNA/RNA/miRNA Universal Kit (QIAGEN) and following the manufacturer’s protocol, including DNase treatment option. Library preparation was performed using the NEBNext® rRNA Depletion Kit v2 (Human/Mouse/Rat) (New England Biolabs) following the manufacturer’s protocol (including RNase H and DNase I digestions). 100-bp paired-end sequencing was performed on the Illumina HiSeq 4000 or NovaSeq 6000 platform.

### Histone acid extraction, histone derivatization, and analysis of post-translational modifications by nano-LC-MS

4 million cells from each clonal mMSC line were collected and frozen at −80℃. Thawed pellets were lysed in nuclear isolation buffer (15 mM Tris pH 7.5, 60 mM KCl, 15 mM NaCl, 5 mM MgCl2, 1 mM CaCl2, 250 mM sucrose, 10 mM sodium butyrate, 0.1% (v/v) beta-mercaptoethanol, commercial phosphatase and protease inhibitor cocktail tablets) containing 0.3% NP-40 alternative on ice for 5 min. Nuclei were subsequently washed twice in the same buffer without NP-40, and pellets were resuspended using gentle vortexing in chilled 0.4 N H2SO4, followed by a 3 hr incubation while rotating at 4℃. After centrifugation, supernatants were collected and proteins were precipitated in 20% TCA overnight at 4℃, washed with 0.1% HCl (v/v) acetone once, followed by two washes with acetone alone. Histones were resuspended in deionized water. Acid-extracted histones (20 μg) were resuspended in 50 mM ammonium bicarbonate (pH 8.0), derivatized using propionic anhydride and digested with trypsin as previously described (15). After the second round of propionylation, the resulting histone peptides were desalted using C18 Stage Tips, dried using a centrifugal evaporator and reconstituted using 0.1% formic acid in preparation for LC-MS analysis. Nanoflow liquid chromatography was performed using a Thermo Fisher Scientific, Vanquish Neo UHPLC equipped with an Easy-Spray™ PepMap™ Neo nano-column (2 µm, C18, 75 µm X 150 mm). Buffer A was 0.1% formic acid and Buffer B was 0.1% formic acid in 80% acetonitrile. Peptides were resolved using at room temperature with a mobile phase consisting of a linear gradient from 1 to 45% solvent B (0.1% formic acid in 100% acetonitrile) in solvent A (0.1% formic acid in water) over 85 mins and then 45 to 98% solvent B over 5 mins at a flow rate of 300 nL/min. The HPLC was coupled online to an Orbitrap Exploris 240 (Thermo Scientific) mass spectrometer operating in the positive mode using a Nanospray Flex Ion Source (Thermo Fisher Scientific) at 1.9 kV. A full MS scan was acquired in the Orbitrap mass analyzer across 350–1050 m/z at a resolution of 120,000 in positive profile mode with an auto maximum injection time and an AGC target of 300%. Parallel reaction monitoring experiments were followed for monitoring the targeted peptides based on the inclusion list. Targeted ions were fragmented using HCD fragmentation. These scans typically used an NCE of 30, an AGC target standard, and an auto maximum injection time. Raw files were analyzed using EpiProfile 2.0^54^ and Skyline^55^. The area for each modification state of a peptide was normalized against the total signal for that peptide to give the relative abundance of the histone modification. Calculated MS ratios (Supplementary Table 2)

### Whole genome bisulfite sequencing

Whole genome sequencing libraries were generated from 1000 ng of genomic DNA extracted from approximately 4 million cells and fragmented to 300-400 bp using the Covaris focused-ultrasonicator E210. The NxSeq AmpFREE Low DNA Library Kit (Lucigen - CA10837-144) was then used. End repair, A-tailing, adapter ligation, and clean-up steps were performed as per the manufacturer’s recommendations. Bisulfite conversion was performed using the EZ-96 DNA Methylation-Gold™ MagPrep Kit (Zymo Research) according to the manufacturer’s protocol. Bisulfite-converted DNA was amplified by PCR using the KAPA Hifi HotStart Uracil+ Kit (Roche) according to the manufacturer’s protocol. The amplified libraries were then purified using AMPure XP Beads (Beckman Coulter). 150-bp paired-end sequencing of the WGBS libraries was performed using the Illumina HiSeqX system.

### Hi-C sample preparation

C3H10T1/2 mouse mesenchymal stem cells (two million cells per sample) were crosslinked and processed using the Arima-HiC+ kit and following the Arima Genomics HiC protocol for Mammalian Cell Lines (A510008). Proximally-ligated DNA was fragmented to an average size of 400 bp using the BioRuptor UCS-300 at low intensity for 10 cycles (30s ON, 90s OFF). Library preparation of fragmented proximally-ligated DNA was performed using the NEBNext® Ultra™ II DNA Library Prep Kit (E7645S), NEBNext Multiplex Oligos for Illumina (E6440S), and the Arima HiC+ Kit according to manufacturer protocols. Hi-C libraries were sequenced by the McGill University Genome Centre at 150 bp paired-end reads using the Illumina NovaSeq 6000.

### Immunofluorescent staining and analysis

Cells were seeded on glass coverslips until between 60-80% confluent and fixed with a cold solution of methanol:acetone at 80:20 ratio for 15 minutes. The samples were blocked 1h with 5% skim milk followed by 1h with rabbit anti-H3K9me3 antibody (Supplementary Table 1) at 1:400 dilution and 1h with Alexa Fluor-488 goat anti-rabbit secondary antibody, all in blocking solution. Samples were then counterstained with 4′,6-diamidino-2-phenylindole (DAPI) solution in PBS for 5 min. All the different treatments were separated by 3 washes with PBS. The coverslips were inverted onto permafluor mountant (epredia) on microscope slides, viewed and imaged on confocal LSM800 at the McGill University Advanced BioImaging Facility (ABIF). Z-stacks of 10 slices were taken and image projection (average setting) was built using Fiji^56^ v.2.15.1. All images were taken under the same conditions, but laser strength and exposure were slightly adjusted for each image so that there is no saturation and ensuring similar fluorescence intensity in each image. This allowed a proper comparison of the spatial distribution of the foci. Images were filtered using ImageJ^57^ v.1.54 for any nuclei that were dividing and any nuclei that were cutoff at the border of the images. Analysis was performed using the open-source software CellProfiler^58^ v.4.2.6. A CellProfiler pipeline was designed to detect and count nuclei as well as the number of H3K9me3 foci within each nucleus using the same parameters for all images. For detecting nuclei, a manual thresholding method was applied with a threshold of 0.015 and only nuclei with a diameter from 50 to 350 included. 130 nuclei per cell line was included for analysis. For detecting the number of H3K9me3 foci within each nucleus, a manual thresholding method was applied with a threshold of 0.1, smoothing filter of 11 and minimum allowed distance of 6 (which distinguishes between clumped foci). The number of foci per nucleus in the QuiKO cells were normalized by multiplying the foci count per nucleus by the ratio of the mean surface area of the PA nuclei to the mean surface area of the QuiKO nuclei. Furthermore, the average distance of foci to the center of each nucleus was normalized by dividing by the radius.

### Visualization

Unless otherwise stated, figures were created using ggplot2^59^ v3.3.0. Coverage/alignment tracks were visualized using pyGenomeTracks^60^ v3.2.1 or Trackplot^61^ v.1.5.10. Heatmaps and aggregate (average) profiles of bigwig pileups at enhancers, strong enhancers, promoters, gene bodies and other ATAC-seq peaks sets were generated using the computeMatrix, plotProfile and plotHeatmap functions from deepTools^62^ v3.3.1. For all of these heatmaps and aggregate profiles, replicates (n=2 or n=3) were merged prior to plotting.

### ChIP-seq, CUT&RUN, ATAC-seq and RNA-seq processing and analysis

ChIP-seq reads were processed as described^2^, where they were aligned using BWA^63^ v.0.7.17 to a combined reference of mm10 and dm6 and afterwards filtered using a cut-off of MAPQ < 3 using Samtools^64^. For Cal27-HNSCC, ChIP-seq reads were aligned using a combined reference of hg38 and dm6. Samclip (https://github.com/tseemann/samclip) v.0.2 (samtools view -h in.bam | samclip --ref ref.fa | samtools sort > out.bam) was used to filter bacterial sequences prior to alignment for all ChIP-seq reads.

CUT&RUN reads were trimmed of adapter sequences using Fastp^65^ v0.23.3. Bowtie^66^ v2.5.1 was used to align the reads to the mm10 assembly using the parameters “-I 10 -X 700 --end-to-end --local --very-sensitive-local --no-unal --no-overlap --no-dovetail --no-mixed --no-discordant -1 ${SAMPLE}.R1.fq.gz -2 ${SAMPLE}.R2.fq.gz”. Samtools was used to remove duplicated reads and reads below a cutoff of MAPQ < 3. Samclip was used to filter bacterial sequences prior to alignment.

The bamCoverage function from deepTools was used to normalize ChIP and CUT&RUN signals by dividing by the total alignments (in millions) (bamCoverage -b $BAM -o $OUTPUT.bigWig --normalizeUsing CPM --centerReads -bs 10 -e 200 -bl $mm10.BLACKLIST.bed). To generate coverage tracks, following CPM (counts per million) normalization as indicated above, replicates (n=3) were merged in a stepwise fashion using bigwigCompare from deepTools with parameters ‘-b1 rep1 -b2 rep2 $outdir --operation mean -bs 10 -o $merged.step1.cpm.bw’ and ‘-b1 merged.step1.cpm.bw -b2 rep3 $outdir --operation mean -bs 10 -o $merged.final.cpm.bw’. A normalization factor was computed by multiplying the genome-wide modification percentage values obtained from mass spectrometry (these values were averaged per condition) by the total number of bins and dividing by the total signal (in CPM) for a given bigWig file, which consists of merged replicates. This normalization factor was then multiplied to the depth-normalized signal (in CPM) for each merged bigWig track to generate MS-normalized coverage tracks. Finally, ChIP-seq bins greater than 100 were set to 100. The ENCODE blacklist ^67^ was used for filtering.

For ATAC-seq, raw reads were trimmed and filtered for quality using Trimmomatic^68^ v0.39. Trimmed reads were mapped to the mm10 genome assembly using Bowtie2, and non-uniquely mapping reads were removed. Afterwards, the reads were adjusted by shifting all positive-strand reads 4 bp downstream and all negative-strand reads 5 bp upstream to the center of the reads on the transposase binding event. Peak calling was performed on each replicate using MACS2^69^ v2.2.6 with ‘-extsize 200 -shift - 100 -nomodel’ parameters. To find a set of reproducible peaks across replicates, we calculated the irreproducible discovery rate (IDR)^70^ and excluded peaks with an IDR greater than 0.05 across every pair of replicates. Subsequently, the ENCODE blacklist was used to filter the peaks. Coverage tracks were generated using bamCoverage with parameters ‘-b $BAM -o $OUTPUT.bigWig --normalizeUsing CPM --centerReads -e 200 --minMappingQuality 5’’. Replicates (n=2) were merged using bigwigCompare from deepTools for aggregate plots.

To generate accessible enhancers for Fig. 1, an intersection of ATAC-seq peaks between TKO and QKO were first generated. From this set of ATAC-seq peaks, peaks within +/- 3kb of TSSs were excluded to account for promoter biases - these filtered ATAC-seq peaks were then designated as accessible enhancers. Accessible promoters were then defined as ATAC-seq peaks overlapping protein-coding promoters, which were defined as 2-kb regions centered at the TSS (1500 bp upstream and 500 bp downstream). Strong enhancers were called from TKO ChIP-seq H3K27ac using ROSE^37^ v1.0.0 with “-s 12500 -t 2500”, to ensure that H3K27ac peaks within 12.5 kb were stitched together and regions within 2.5 kb of TSSs were excluded. Nearest enhancers, as indicated in Fig. S1g, were derived using the closestBed function from BEDTools^71^ v.2.31.1.

For RNA-seq, raw reads were aligned to mm10 genome build using STAR^72^ v2.5.3a. Afterwards, featureCounts^73^ v1.5.3 was used to count exonic reads from the GTF transcript annotation (GENCODE version from UCSC). DESeq2^74^ v1.26.0 was used to perform differential gene expression analyses. Adjusted log fold changes (LFC) were calculated using ‘apeglm’^75^ v.1.8.0. Significantly differentially expressed genes were selected based on absolute log2FC > 2 and adjusted p-value less than 0.05. Coverage bigWig tracks for RNA-seq signal was computed using bamCoverage from deepTools with the parameters “bamCoverage -b $SAMPLE.bam -o $SAMPLE.RPKM.bw -bl $BLACKLIST.bed --normalizeUsing RPKM”.

To find large H3K36me2 peaks marking clusters of enhancers in TKO, enhancers from Ensembl (www.ensembl.org) and FANTOM^76^ were firstly pooled together for the mm10 assembly. Epic2^77^ v0.0.52 was used to call peaks on two replicates of ChIP-seq H3K36me2 for the TKO cell line and their intersection was taken using BEDTools - these merged peaks were subsequently used to to generate a read count matrix of ChIP-seq TKO H3K36me2 triplicates (n=3) using featureCounts. These peaks were then filtered for having greater than 100 average normalized read count across TKO replicates and overlapping at least one annotated enhancer. Afterwards, the peaks were extended by 50000 bp on each side and these regions were designated as large H3K36me2 enhancer peaks. Next, to generate two groups of genes of similar expression, but differ in whether or not they are located within these large H3K36me2 enhancer peaks, we filtered for protein-coding genes with at least 1 FPKM gene expression at baseline (in the parental sample). Afterwards, the genes were stratified into bins, with each bin grouping genes of similar expression (+/- 3 FPKM). We selected genes with 9-12 FPKM to generate the two groups of genes in Fig. 1f with equal group sizes (n=219 per group): one group of genes within the large H3K36me2 peaks and a randomized, control set of genes outside of these large H3K36me2 peaks.

Genome-wide Pearson’s correlation analysis was performed using the multiBigWigSummary bins program from DeepTools using 500 bp bins. The results were plotted using gplots (https://github.com/cran/gplots) v3.1.3.1 ‘heatmap.2’ function as heatmaps.

For Fig. 2, genes marked with residual H3K36me2 were determined by taking genes overlapping QKO H3K36me2 peaks by at least 1bp. These remaining 119 QKO H3K36me2 peaks were generated as described previously^22^. To generate the correlation scatter plot between the size of H3K36me2 peak and gene expression fold changes, featureCounts was used to generate a count matrix from QKO ChIP-seq H3K36me2 replicates (n=3), using the remaining 119 peaks in QKO as reference. Afterwards, the raw counts per replicate were normalized by dividing by their total alignments (in millions) to generate counts per million (CPM) and the average was taken amongst the replicates for each of the 119 peaks. Afterwards, BEDtools was used to find the overlapping gene for each of the 119 peaks, and subsequently the gene/peak pairs were plotted using R. To generate accessible enhancers for Fig. 2, an intersection of ATAC-seq peaks between QKO and QuiKO were first generated. From this set of ATAC-seq peaks, peaks within +/- 3kb of TSSs were excluded to account for promoter biases - these filtered ATAC-seq peaks were then designated as accessible enhancers. Next, we filtered for accessible enhancers overlapping the 119 remaining H3K36me2 peaks found in QKO. Furthermore, ATAC-seq peaks that overlapped with promoters that were marked by residual QKO H3K36me2 peaks were designated as accessible promoters for Fig. 2. Accessible enhancers and promoters devoid of H3K36me2 were ATAC-seq peaks that did not overlap the 119 remaining H3K36me2 peaks in QKO.

To generate metagene plots, 21833 protein-coding genes (from Ensembl) were selected with gene length >= 25th percentile (6842 bp) and <= 75th percentile (48614 bp) to avoid over/under-scaling gene sizes when generating pileups for the aggregate plots. Furthermore, adjacent/neighboring genes within −2 kb TSS and +2 kb TES regions were removed to eliminate edge effects. From this set of filtered protein-coding genes, genes with zero normalized counts were designated as “silent genes” (n=2000). Lastly, to generate the expressed gene groups, protein-coding genes below 0.1 FPKM in the parental cell line were removed and from the remaining genes, the bottom 2000 genes designated as lowly expressed genes whereas the top 2000 genes were designated as highly expressed genes. These three gene groups (n=2000 per group) were then used as reference to generate metagene plots using computeMatrix from deepTools with the parameters “computeMatrix scale-regions -R $SILENT_GENES.bed $LOWLY_EXPRESSED.bed $HIGHLY_EXPRESSED.bed -S $SAMP.bw -bl $BLACKLIST.bed -b 2000 -a 2000 -m 20000 -bs 500 --missingDataAsZero --skipZeros” and plotted using plotProfile from deepTools. Genes with significant H3K27me3 or H3K9me3 in their gene bodies were selected (adjusted p-value < 0.05) from a pool of mm10 protein-coding genes, in which featureCounts was used on aligned ChIP-seq files with mm10 protein-coding genes as the annotation/reference followed by DESeq2 to generate differentially-marked (either by H3K27me3 or H3K9me3 respectively) protein-coding genes.

“Peakiness” scores were computed as the average read-depth normalized coverage of the top 1% most covered 1-kb windows across the genome, excluding those overlapping blacklisted regions, as previously described^2^.

To generate genome-wide binned analysis in Fig. 5, raw H3K9me3 ChIP-seq tag counts were firstly binned into 100 kb windows using BEDtools with intersectBed (-c) in combination with the makewindows command, as previously described^18^. Binned tag counts were library-sized normalized by dividing by the total number of mapped reads after filtering and subsequently input-normalized by taking the log2 ratio of ChIP signals by those of the input (i.e., without immunoprecipitation), with the addition of a pseudocount (1e-15) to avoid division by 0: log2(((H3K9me3 ChIP binned tag counts)/(H3K9me3 total ChIP mapped reads)+1e-15)/((input binned tag counts)/(total input mapped reads)+1e-15)). Additionally, quantitative normalization was performed by multiplying the signal (log2 ratio over input as described above) by the genome-wide modification percentage information obtained from mass spectrometry. Genic and intergenic regions were annotated as described previously^18^. Similarly behaving bin clusters were identified using HDBSCAN^78^ v0.8.24 with the same parameters for all comparisons (minPts = 1000, eps = 1000). To better reflect the large megabase-sized domains of H3K9me3, 100-kb bins within 1-mb were merged together for each bin cluster. Afterwards, regions in cluster B overlapping with regions in cluster A were excluded.

Transposable elements (TEs) were obtained from RepeatMasker (www.repeatmasker.org). These TEs were used as the annotated reference file for featureCounts to count RNA-seq transcript expression and subsequently generate a count matrix for input into DESeq2 to compute differentially expressed TEs. Adjusted LFC were calculated using ‘apeglm’ and significantly differentially expressed TEs were selected based on absolute log2FC > 1 and adjusted p-value less than 0.05.

### WGBS processing and analysis

Raw WGBS reads were processed and analyzed as previously described^2^, where they were mapped and methylation calling was performed using Bismark^79^ against the mouse assembly mm10, ENCODE blacklisted regions were removed and only CpGs covered by ≥5× in all samples were retained for the computation of DNA methylation levels.

### Hi-C processing and analysis

Raw Hi-C reads were trimmed of adapter sequences using Fastp and aligned to the mm10 assembly using BWA with default parameters. Afterwards, ligation junctions (Hi-C pairs) were identified in aligned paired-end sequences using Pairtools (https://pairtools.readthedocs.io/en/latest/) v1.1.0. The restriction enzyme specified to Pairtools was the Arima kit, which uses the two restriction enzymes: DpnII and HinfI. Duplicated reads were marked using ‘pairtools dedup’ with the parameters: “--mark-dups --backend cython --output-stats $SAMPLE.stats -o $SAMPLE.dedup.pairs.gz”. Then, duplicated reads were removed and only uniquely-mapped reads were selected using ‘pairtools select’ with the parameters: “--chrom-subset mm10.canon.chroms ’(rfrag1!=rfrag2) and ((pair_type==“UU” or pair_type==“RU” or pair_type==“UR”))’ -o $SAMPLE.pairs.gz ${samp}.dedup.pairs.gz’. Junctions were subsequently binned and counted using Cooler^80^ v0.10.0, in which pairs (.pairs.gz) files were converted to a cooler (.cool) file using “cooler cload pairix --assembly mm10 mm10.canon.sizes:1000 $SAMPLE.pairs.gz $SAMPLE.1000.cool”. The cooler file underwent matrix balancing using “cooler balance $SAMPLE.1000.cool --blacklist $BLACKLIST.bed”. The matrix was then binned into different window sizes contained within a single multi-resolution cooler (.mcool) file using “cooler zoomify --out $SAMPLE.mcool --resolutions 1000,5000,10000,25000,50000,100000,250000,500000,1000000,2500000 --balance --balance-args --max-iters 1000 $SAMPLE.1000.cool”.

Cooltools^81^ v0.4.0 was used to compute compartment scores (eigenvalues): “cooltools eigs-cis --phasing-track mm10.100000.gc $SAMPLE.mcool::/resolutions/100000”, with the binned profile of GC content (mm10.100000.gc) as the reference track for sign flipping. To assess interaction patterns between bins belonging to different quantiles of compartment scores, the compute-saddle function from cooltools with default parameters at 100-kb resolution was implemented, using eigenvectors and expected values from cooltools’ eigs-cis (calling compartments) and compute-expected modules, respectively. To quantify the interaction strength between A-A and B-B compartments, we took the top 20% of B-B interactions and the top 20% of A-A interactions, normalized by the bottom 20% of A-B interactions, as previously described^82^. Long-range *cis* contacts were defined as interactions spanning greater than 10 megabytes (mb) and short-range *cis* contacts were defined as interactions 1-kb to 10-mb.

Insulation scores were computed for ‘.mcool’ files at 10-kb resolution with a window size of 100kb using the “cooltools insulation” module with default parameters. Only boundaries annotated as “is_boundary_100000 == “TRUE”” were included. Loop scores were computed using the “cooltools dots” module with default parameters at 25-kb resolution. Long-range loops were defined as those spanning greater than 500-kb whereas short-range loops were defined as those spanning 500-kb or less.

### Statistical analyses

The statistical details used in each analysis are provided in the figure legends wherever applicable. Wilcoxon’s rank-signed sum test was used in all statistical comparisons. Differences were considered significant for *p-*value less than 0.05. Boxplots were generated using R. In the box plots, boxes span the lower (first quartile) and upper quartiles (third quartile), median is indicated with a center line and whiskers extend to a maximum of 1.5 times the interquartile range. For barplots, error bars display the standard deviation around the mean. Results for correlation analyses are reported for Pearson’s correlation coefficient (*R*) and the associated *p*-value where applicable.

### Data availability

Previously published and publicly available ChIP-seq data can be accessed via the National Center for Biotechnology Information Gene Expression Omnibus (NCBI-GEO): H3K9me3 (parental) under accession number GSE118785, H3K27me3 (parental, SETD2-KO, H3K36M, and NSD1-2-DKO samples) under accession number GSE160266, and H3K36me2 (parental, ASH1L-KO, NSD3-KO, NSD1-2-SETD2-TKO, NSD1-2-3-SETD2-QKO, and NSD1-2-3-SETD2-ASH1L-QuiKO samples) under accession number GSE243566. Newly generated ChIP-seq, ATAC-seq, CUT&RUN, WGBS, RNA-seq and Hi-C data can also be accessed under accession number GSE274367.

### Code availability

The source code for bioinformatic analysis is available on request.

## Additional files

Supplementary Table 1 (.xlsx): Antibody list

Supplementary Table 2 (.xlsx): Calculated mass-spectrometry values

## Acknowledgements

We would like to thank all members of the J.M. lab for their input and feedback. We thank the McGill University Genome Centre for their expertise in sample quality verification and sequencing services. We thank Dr. Daniel Weinberg and the lab of Dr. Chao Lu for earlier work that generated many of the resources (C3H10T1/2 NSD1/2-DKO, NSD1/2-SETD2-TKO, and H3.3K36M-OE cell lines) that became the core of this research. We also thank the group of Dr. Benjamin Garcia for their expertise in mass spectrometry sample processing.

## Authors’ contributions

R.P. and G.S. contributed equally to this work. Study design, G.S., R.P., and J.M. Writing (original draft), G.S., R.P., and J.M. Laboratory experiments, G.S and C.H. Bioinformatic analyses, R.P. Data processing, R.P. and E.B. Supervision, J.M. Funding acquisition, J.M. All author(s) read and approved the final manuscript.

